# The Cappuccino interactome reveals an intracellular role for Semaphorin-2a in *Drosophila* oogenesis

**DOI:** 10.64898/2026.05.11.724452

**Authors:** Carolyn Wu, Sudeepa Rajan, Merin Rixen, James Wohlschlegel, Margot Elizabeth Quinlan

## Abstract

The spatiotemporal regulation of an actin mesh during *Drosophila* oogenesis is essential for proper localization of cell polarity determinants that establish the future patterning of the embryo. Here, we reveal an unexpected role for Semaphorin-2a (Sema2a) in actin mesh regulation and oogenesis. Sema2a classically functions as a secreted guidance cue that binds its cognate Plexin-B (PlexB) receptor to establish neural circuits. In contrast, we find that Sema2a is expressed inside the germarium, germline, and follicle cells of the developing ovary. Sema2a mutants possess small ovaries that fail to develop past mid-oogenesis. We demonstrate that Sema2a interacts with Cappuccino (Capu), a key actin nucleator crucial for building the actin mesh in *Drosophila* oocytes. Sema2a inhibits the actin assembly activity of Capu *in vitro*. Furthermore, genetic interaction between Sema2a and Capu influences mesh density and disrupts *oskar* mRNA localization. PlexB mutants, however, exhibit wild-type size ovaries with *osk*ar mRNA localization distinct from Sema2a mutants, confirming the non-canonical role of Sema2a in oogenesis.

**Summary:** This study reveals a novel interaction between the actin nucleator Cappuccino and the typically secreted neural guidance factor Semaphorin-2a. It is shown that Semaphorin-2a inhibits the actin polymerization activity of Cappuccino *in vitro* and play an intracellular role in oogenesis.

## Introduction

Proper regulation of actin is crucial for egg development. For example, an isotropic actin mesh that fills the *Drosophila* oocyte during mid-oogenesis is critical to microtubule organization and polarity establishment (Dahlgaard et al., 2007). The mesh slows microtubule-driven cytoplasmic flows, facilitating proper transport of several mRNAs, including *oskar* (*osk), bicoid (bcd), gurken (grk), and nanos (nos),* that define the anterior-posterior and dorsal-ventral axes of the oocyte and future embryo. By late-oogenesis, the mesh disappears, and fast, microtubule-dependent streaming ensues (Dahlgaard et al., 2007). Disappearance of this mesh controls the transition from slow to fast streaming, during which cytoplasmic flows are approximately 10 times faster than slow streaming to allow efficient distribution of mRNAs and proteins as the oocyte increases in size (Quinlan, 2016). Understanding the molecular machinery and mechanisms underlying the dynamically regulated actin mesh is important because absence of the mesh and premature fast streaming lead to loss of polarity and infertility (Dahlgaard et al., 2007; Manseau and Schupbach, 1989; Theurkauf, 1994), while persistence of the mesh prevents fast streaming and delays mRNA delivery, decreasing fertility (Bor et al., 2015; Dahlgaard et al., 2007).

A genetic screen first identified *cappuccino* (*capu)* and *spire (spir)* as maternal-effect loci responsible for patterning the anterior-posterior and dorsal-ventral axes of the embryo (Manseau and Schupbach, 1989). These two genes were implicated in regulating the cytoskeleton, as mutations in both *capu* and *spir* resulted in premature or delayed streaming, leading to disruption of *osk, grk,* and *nos* localization (Bor et al., 2015; Emmons et al., 1995; Manseau et al., 1996; Manseau and Schupbach, 1989; Theurkauf, 1994; Wellington et al., 1999). Capu is part of the FMN family of formin nucleators and Spir is the defining member of a class of actin nucleators commonly referred to as tandem WASP homology 2 (WH2) domain nucleators (Dominguez, 2016; Higgs and Peterson, 2005). They both nucleate actin, but Capu also processively elongates actin filaments. Deeper investigation into the roles of Capu and Spir led to the finding that these two proteins build the actin mesh (Dahlgaard et al., 2007; Quinlan, 2013). Functional interaction between Capu and Spir is observed both *in vitro* and *in vivo* (Bradley et al., 2019; Montaville et al., 2014; Quinlan, 2013; Quinlan et al., 2007; Vizcarra et al., 2011; Zeth et al., 2011). In *Drosophila,* point mutations that prevent Capu/Spir interaction result in failure to build an actin mesh, premature fast streaming, and fertility loss (Quinlan, 2013; Vizcarra et al., 2011). Moreover, during late-oogenesis, Capu and Spir protein levels decrease to less than 10% of those in the whole ovary (Quinlan, 2013). The reduction in protein levels coincides with the disappearance of the actin mesh, suggesting that their activity needs to be regulated for proper mesh formation and removal. An analogous mesh is also found in mouse oocytes, where mammalian homologs of Capu and Spir, Formin-2 (Fmn2) and Spire1/Spire2 respectively, are required to build an actin network for asymmetric spindle positioning (Azoury et al., 2011; Pfender et al., 2011; Schuh, 2011). Thus, the importance of Capu/Spir is evolutionarily conserved, yet there is little known about the factors that regulate these proteins or the actin mesh.

Large-scale proteomic screens are powerful tools for studying protein-protein interactions in a range of biological processes. Conventional methods like co-immunoprecipitation (co-IP) require stable, high-affinity complexes to survive cell lysis while more current methods like proximity labeling allow detection of transient interactions in a native environment. Combining these complementary methods offers a more complete view of the protein interaction landscape and provides added confidence when novel interactors are enriched in both datasets. With the goal of understanding Capu and Spir as well as actin mesh regulation, we combined co-IP and proximity labeling with quantitative mass spectrometry to identify both transient and stable interactors and build a comprehensive map of the Capu/Spir interactome. We used GFP-Traps for co-immunoprecipitation and the biotin ligase, TurboID, to perform proximity labeling (Baker et al., 2021; Branon et al., 2018). These two approaches validate known Capu and Spir interactors as well as reveal previously unrecognized interactors.

Here, we elucidate the unanticipated, intracellular role of the novel Capu interactor, Semaphorin-2a (Sema2a). Semaphorins and their cognate plexin receptors are known for their role in neural guidance. Indeed, Sema2a has been extensively studied in the central nervous system (CNS) as a secreted signaling protein that binds to its canonical receptor, Plexin-B (PlexB), to mediate axon guidance (Ayoob et al., 2006; Hu et al., 2001; Roh et al., 2016). In fact, the large family of Semaphorin signaling proteins contributes to a range of processes in development (Jongbloets and Pasterkamp, 2014). Notably, Sema2a mRNA is detected in the embryonic gonad, in addition to neurons, suggesting it could play a role in gametogenesis (Kolodkin et al., 1993). Here we describe a role for Sema2a in oogneesis. Although normally secreted, we observe that Sema2a is abundant inside the germarium, germline, and follicle cells of the ovary. Confirming its importance in oogenesis, loss of Sema2a leads to decreased ovary size, with ovaries often too small to find or possibly non-existent. When the ovaries are visible, egg chambers have not developed past mid-oogenesis. Loss of PlexB results in ovaries that develop normally and are wild-type in size, indicating that Sema2a functions independently of PlexB in oogenesis. Using biochemical assays and genetic analyses, we demonstrate that Sema2a interacts with Capu. In fact, Sema2a inhibits the actin assembly activity of Capu *in vitro*. Consistently, we find that actin mesh density as well as *osk* mRNA localization are sensitive to relative levels of Sema2a and Capu. While *osk* mRNA localization is also altered in PlexB mutants, the phenotypes are distinct. Together, our findings establish a previously uncharacterized, intracellular role for Sema2a in oogenesis that is independent of PlexB and provide new insight into the regulation of the actin cytoskeleton during oogenesis.

## Results

### The Capu/Spir interactome identified with quantitative mass spectrometry

To identify the interactors of Capu and Spir, we performed co-immunoprecipitation combined with mass spectrometry (co-IP/MS) in ovaries expressing UASp-GFP-CapuA, UASp-CapuA-GFP, or UASp-SpirC-GFP driven by *capu*-GAL4, a “Trojan-GAL4” driver within the *capu* locus and optimized for germline expression (Figure 1A)(Bailey et al., 2024; Diao et al., 2015). We compared these data to the results of co-IP/MS data from wild-type flies and negative controls lacking the driver. Data-independent acquisition (DIA) mass spectrometry data processed by DIA-NN and analyzed by FragPipe Analyst (FC > 2 and p-value < 0.05) revealed 9 significantly enriched proteins for GFP-tagged Capu and 130 proteins for GFP-tagged Spir, in three biological replicates (Figures 1B-E and Table 1) (Demichev et al., 2020; Hsiao et al., 2024). As expected, Capu and Spir were both significantly enriched in their respective co-IPs. In order to distinguish potential interactions through actin filaments, we repeated the co-IPs in the presence of the actin depolymerizing agent, Latrunculin A. The same sets of proteins were enriched, indicating that none of the interactions that we report here are solely indirect binding to Capu or Spir through filamentous actin (Figures S1A-B). We performed GO analysis using clusterProfiler in R and found that for Spir co-IPs, the top biological processes compared to the background proteome were related to actin and mRNA localization, which are consistent with known functions of Spir (Figure 1G) (Yu et al., 2012). Interestingly, for Capu co-IPs, the top biological processes were more specifically locomotion- and axon-related, based on the genes w, Sema2a, Sema2b, and homer (Figure 1F, Table 1).

**Figure 1.**
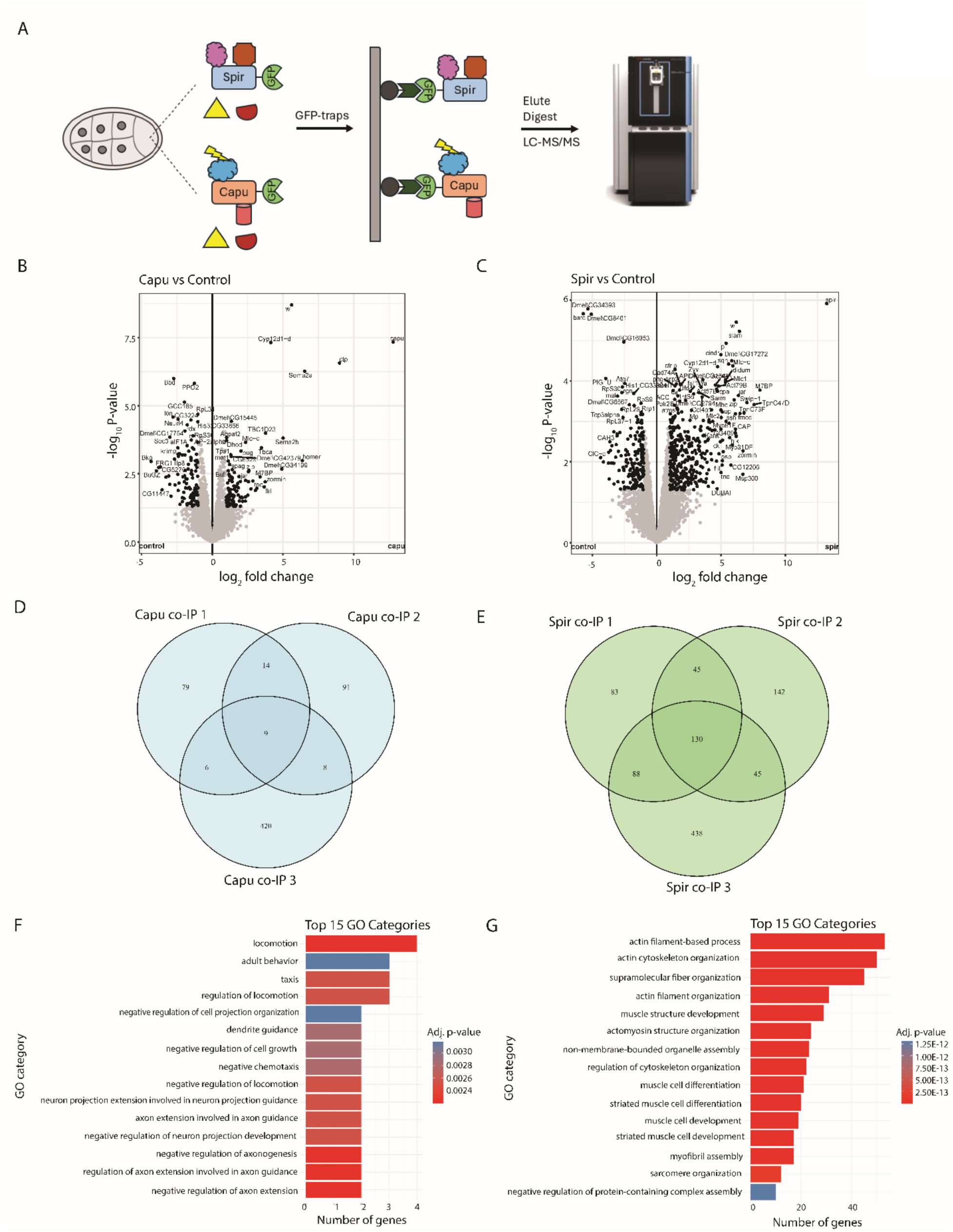
Characterization of Spir and Capu interactomes using co-IP/MS. (A) Schematic showing the experimental workflow. GFP-tagged SpirC and CapuA from lysed ovaries were incubated with ChromoTek GFP-trap magnetic agarose beads. Proteins bound to beads were eluted, digested, and analyzed on an Orbitrap Astral mass spectrometer. Three biological replicates were tested in triplicate for each construct, SpirC-GFP, CapuA-GFP, GFP-CapuA. (B-C) Volcano plots of Capu or Spir compared to controls. Gene names shown represent those that have FC > 2 and p-value < 0.05. (D-E) Venn diagrams comparing common proteins shared across three independent replicates of Capu or Spir co-IPs. (F-G) Plot from clusterProfiler in R, showing enriched annotated biological processes of Capu and Spir co-IPs.

**Table 1.**
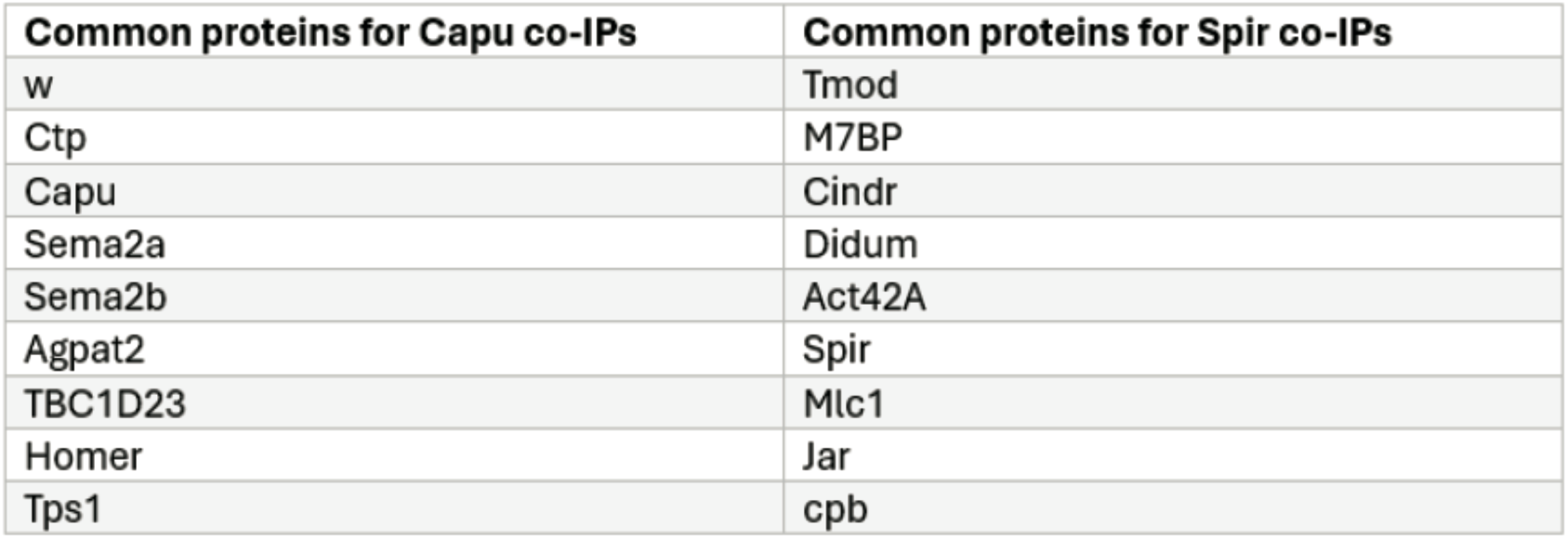
Proteins enriched across all 3 replicates for Capu and Spir co-IPs.

| **Common proteins for Capu co-IPs** | **Common proteins for Spir co-IPs** |
| --- | --- |
| w | Tmod |
| Ctp | M7BP |
| Capu | Cindr |
| Sema2a | Didum |
| Sema2b | Act42A |
| Agpat2 | Spir |
| TBC1D23 | Mlc1 |
| Homer | Jar |
| Tps1 | cpb |

In parallel, we performed TurboID-proximity labeling experiments in ovaries expressing UASp-CapuA-GFP or UASp-SpirC-GFP and UASp-GBP-TurboID, driven by matα-GAL4 (Figure 2A). The data were compared to the UASp-GBP-TurboID controls. Ovaries were stained with an anti-GFP antibody to visualize GFP-tagged Capu and Spir constructs, phalloidin for actin structures, and streptavidin for proteins biotinylated by GBP-TurboID. The colocalization of the GFP-tagged Capu/Spir and biotinylated proteins suggested successful labeling (Figures S1E-F’’’). However, expression levels of Capu and Spir were relatively low, and streptavidin was detected throughout the egg chamber in experimental and control flies, making the GBP-TurboID negative control essential (Figures S1C-F’’’). We next purified the biotinylated proteins with streptavidin magnetic beads and identified them by DIA-based mass spectrometry. Capu and Spir were preferentially isolated from their respective TurboID lines, showing that GBP-TurboID was efficiently labeling the GFP-tagged Capu and Spir constructs as well as proximal proteins. We identified 6 components in the Capu-TurboID data set that were similarly significantly co-purifying with the Capu co-IPs and 22 proteins significantly co-purifying in both the Spir-TurboID and Spir co-IPs (Figures 2B-E and Table 2). Similar to the co-IPs, GO analysis of the TurboID data showed enrichment of locomotion for Capu and actin-related processes for Spir (Figures 2F-G).

**Figure 2.**
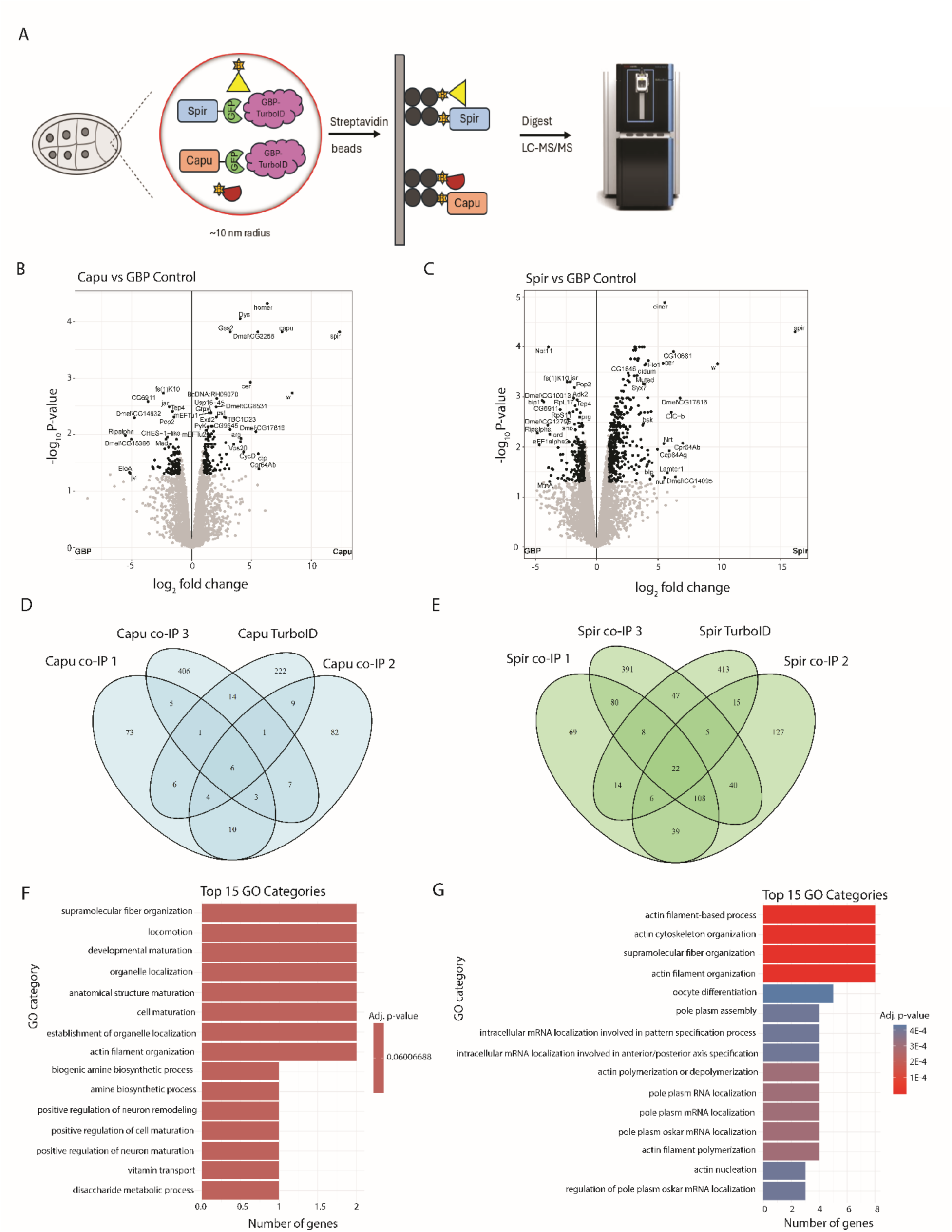
Characterization of Spir and Capu interactomes using TurboID proximity labeling. (A) Schematic showing the experimental workflow. GBP-TurboID biotinylates proteins within ∼10 nm radius of GFP-tagged SpirC and CapuA. Lysate was incubated with streptavidin magnetic beads. Proteins bound to beads were digested and analyzed on an Orbitrap Astral mass spectrometer. Each construct was tested in triplicate. (B-C) Volcano plots of Capu or Spir compared to the controls. Gene names shown represent those that have FC > 2 and p-value < 0.05. (D-E) Venn diagrams comparing common proteins shared across Capu or Spir co-IPs and TurboID. (F-G) Plot from clusterProfiler in R, showing enriched annotated biological processes of Capu or Spir co-IPs and TurboID.

**Table 2.** Proteins enriched across both coIP and TurboID datasets.

| **Common proteins Capu co-IPs and TurboID** | **Common proteins Spir co-IPs and TurboID** |
| --- | --- |
| w | Cindr |
| Ctp | Didum |
| Capu | W |
| TBC1D23 | Arp2 |
| Homer | Cpb |
| Tps1 | Spir |
|  | Ca |
|  | Swip-1 |
|  | cpa |
|  | Tps1 |
|  | Par-1 |
|  | Vinc |
|  | swa |
|  | Dmel\CG11892 |
|  | Strica |
|  | Bor |
|  | Mf |
|  | CLIP-190 |
|  | Arpc4 |
|  | P5CS |
|  | NP_649372 |
|  | Cmb |

Very few Capu or Spir interactors have been identified to date. Supporting the validity of our assays we detected several of the reported binders. For example, we identified Myosin V (*didum*), which interacts with Spir in *Drosophila* and mammalian cells (Figures 1C and 2C) (Ong et al., n.d.; Pylypenko et al., 2016). In the Spir-TurboID dataset specifically, we identified Jun N-terminal kinase (*bsk*), which was previously reported to bind Spir in a yeast two-hybrid assay (Figure 2C) (Otto et al., 2000). We did not expect to detect Capu in Spir datasets because the SpirC isoform we used as bait lacks the KIND domain that binds to Capu. On the other hand, Spir (endogenous, full length) was significantly enriched in the Capu-TurboID dataset, as expected (Figure 2B). Consistent with evidence that the Spir/Capu interaction is transient, we did not observe Spir in Capu co-IP datasets (Figure 1B). Lastly, of the 6 proteins associated with Capu in both co-IP and TurboID datasets, we identified Cut-up (Ctp), which we previously detected with TAP-tagged Capu (Figures 2B, 2D, and Table 2; TAP data not shown). Taken together, these data not only validate biological relevance of the Capu/Spir interactome in actin-related processes, but they also highlight unexpected links between Capu and multiple neuronal proteins.

We were interested in characterizing Sema2a in greater detail considering the extensive work that has been done establishing Sema2a as vital for axon guidance (Ayoob et al., 2006; Hu et al., 2001; Roh et al., 2016). Semaphorins and more specifically, the Semaphorin-Plexin-Mical signaling pathway, play a role in depolymerizing actin networks in growth cones and other motile cells (Ayoob et al., 2006; Fan et al., 1993; Hung et al., 2010; Kolodkin et al., 1993; Luo et al., 1993; Stedden et al., 2019). We hypothesized that Sema2a might interact with Capu to disassemble the actin mesh during oogenesis but the story was more complex. While Sema2a was significantly enriched in the Capu co-IPs (FC ≈ 64, p < 1E-7), we note that it was only moderately enriched in the TurboID dataset (FC ≈ 5, p = 0.0514). Its receptor, PlexB, was also present in the TurboID dataset at even lower levels (FC ≈ 3.5, p = 0.173). Differences in co-IP and TurboID datasets are not uncommon, as each method provides overlapping but also distinct sets of protein based on many factors, such as the duration of a given interaction in the cell (Moreira et al., 2023). The presence of Sema2a at high levels in the co-IPs suggest that Sema2a and Capu bind stably. Its lower enrichment levels in the TurboID experiment suggest that Sema2a may not have accessible lysines in the vicinity of Capu or it may be part of a larger Capu-binding complex but not close enough in proximity to be efficiently labeled.

### Sema2a plays an intracellular role in oogenesis

To test whether Sema2a plays a role in oogenesis, we generated presumptive Sema2a-nulls by crossing the loss-of-function allele *sema2a^03021^*to a deficiency line (Df(2R)B65) *(sema2a^03021^/sema2aDf)*. After eclosion of *sema2a^03021^/sema2aDf* mutants, we observed decreased viability, with an average of ∼50% of flies (N=95 total in 7 replicates) dying upon CO_2_ anesthetization (Figure 3A). An earlier study reported that Sema2a-nulls only survive for 2 days (Kolodkin et al., 1993). We reasoned that the added stress of CO_2_ caused earlier lethality. Very brief CO_2_ exposure resulted in slightly increased viability. Among the flies that survived (n=65 total), around 36% had relatively normal-sized ovaries after overnight fattening with yeast paste. The remaining 64% had small ovaries, if they could be found at all (Figures 3B,C). Despite being fattened, these small ovaries did not contain egg chambers that matured beyond mid-oogenesis, showing the importance of Sema2a in oogenesis (Figure 3C). The ovaries of flies that did not survive CO_2_ exposure had no observable ovaries but could not be directly compared because they could not be fattened. In contrast, *sema2a^03021^/+* and *sema2aDf/+* heterozygotes appeared wild-type, with no loss of viability or diminished ovary-size (Figure 3C).

**Figure 3.**
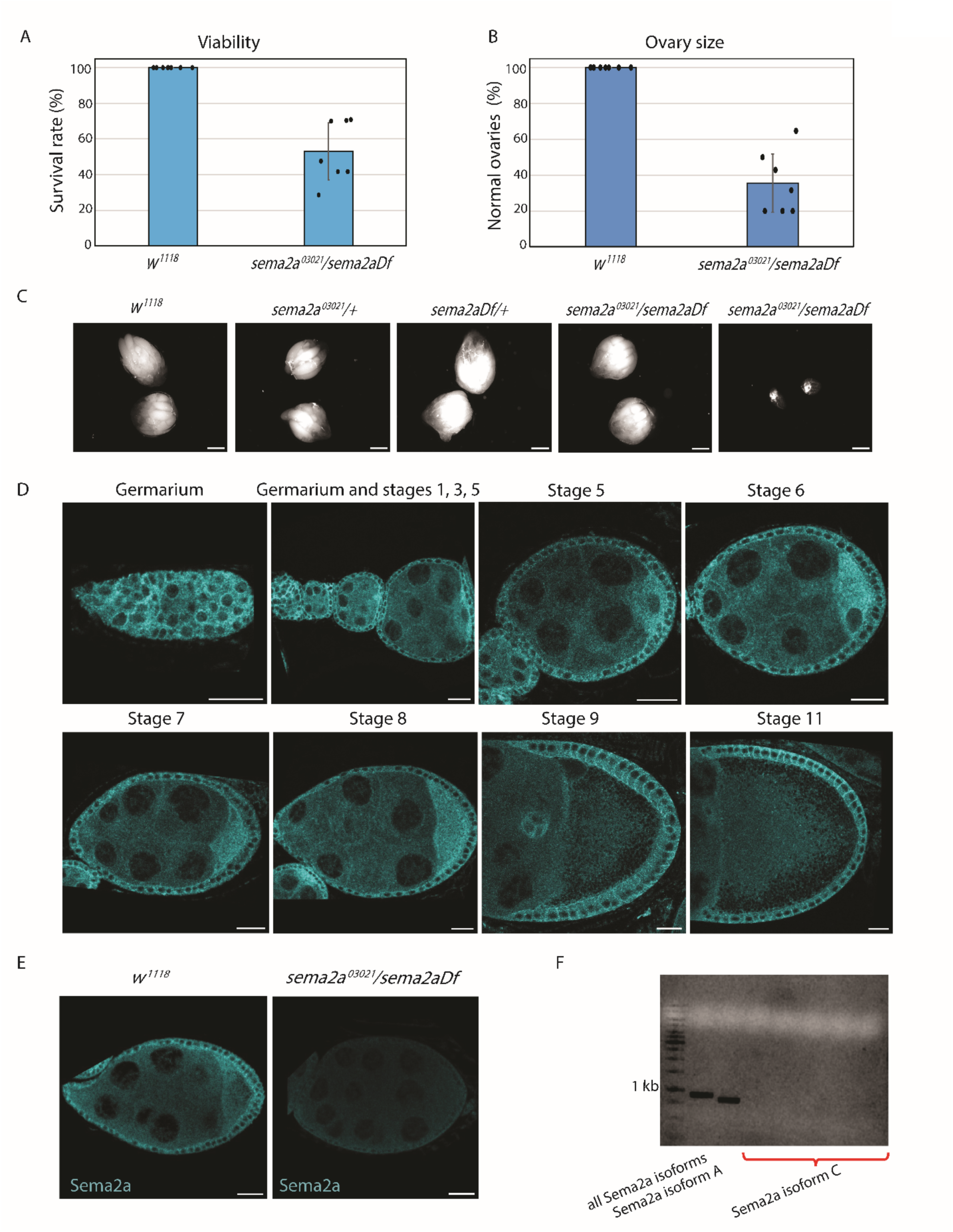
Sema2a plays an intracellular role in oogenesis. (A) Viability of *sema2a*^03021^/*sema2aDf* flies compared to *w^1118^*. Individual dots represent biological replicates. N=95 across all 7 replicates. (B) Ovary size of *sema2a^03021^/sema2aDf* flies compared to *w^1118^*. Individual dots represent replicates from those that survived in (A). (C) Brightfield images of *sema2a^03021^/sema2aDf* ovaries compared to wild type and heterozygotes. Both normal sized (∼36%) and small (∼64%) ovaries are shown for *sema2a^03021^/sema2aDf.* Scale bars are 1 mm. (D) Sema2a detected with anti-Sema2a antibody. The protein is evident in the germarium. It is enriched in the oocytes from ∼stage 3 – stage 9. Sema2a is present in the somatic follicle cells in all stages of the ovariole and in the migrating border cells during stage 9. Scale bars are 20 µm. (E) Sema2a staining for *w^1118^* and *sema2a^03021^/sema2aDf* stage 7 egg chambers. *sema2a^03021^/sema2aDf* shows substantial decrease in signal compared to *w^1118^* and confirms loss of Sema2a. (F) Agarose gel electrophoresis of RT-PCR samples amplifying a sequence present in all isoforms, Sema2a isoform A sequence, or Sema2a isoform C sequence. A total of three pairs of primers detected no product for isoform C. The gel above shows one primer set with a temperature gradient.

Because Sema2a acts in axon guidance as a secreted protein, we asked whether it localizes within the egg chamber. We stained ovaries with an antibody against Sema2a that had been validated in motor axons (Figure 3D) (Winberg et al., 1998). To confirm the specificity of the Sema2a antibody in the egg chamber, we stained *sema2a^03021^/sema2aDf* mutants and found vastly reduced staining compared to the abundant signal in the oocyte and follicle cells of wild-type egg chambers (Figures 3E and S2A). Importantly, Sema2a was highly expressed throughout the ovariole. We detected it in the early germarium, consistent with reports with a GFP-trapped Sema2a (Figure 3D) (Hsu and Drummond-Barbosa, 2017). It also filled early egg chambers, both somatic follicle cells and the germline cyst. By stage 3, Sema2a was enriched in the oocyte compared to the nurse cells. High levels of Sema2a were seen primarily in the oocyte and follicle cells during early and mid-oogenesis. By stage 9, however, Sema2a levels in the oocyte decreased. We detected little to no Sema2a in stage 11 germline cells, though protein remained in the follicle cells (Figure 3D).

We questioned how it is possible that Sema2a localizes at such high levels in the oocyte and follicle cells, given that Sema2a is known to be secreted in the CNS (Bates and Whitington, 2007; Kolodkin et al., 1993; Wu et al., 2011). Upon examination of the Sema2a gene structure on FlyBase (Gramates et al., 2022; Öztürk-Çolak et al., 2024), we found that while Sema2a splice variants A, B, and E have a signal peptide sequence, Sema2a variant C has a distinct N-terminal extension lacking this sequence. Thus, we hypothesized that Sema2a-C was the intracellular variant we were observing. To determine which splice variant(s) were expressed in the egg chamber, we performed reverse transcription polymerase chain reaction (RT-PCR) on RNA extracted from *w*^1118^ (wild-type) ovaries. We reverse transcribed Sema2a mRNA, performed first strand cDNA synthesis, and amplified the N-terminal region specific to the Sema2a isoforms. We first confirmed the presence of Sema2a by amplifying a region present in all isoforms, as indicated by the band at ∼900 bp (Figure 3F). We were unable to detect Sema2a isoform C with multiple primer pairs (Figure 3F). Nor were we able to detect Sema2a isoform E (data not shown). Because the Sema2a isoform B sequence is entirely contained within isoform A, its presence cannot be confirmed or ruled out. However, isoform A can be uniquely identified due to a short, unique sequence not present in isoform B. Sema2a isoform A expression was abundant, based on the prominent band at ∼800 bp (Figure 3F). Sequencing the amplified product corroborated that it was the sequence specific to isoform A (Figure S2B). We conclude that Sema2a isoform A and not isoform C is present in the egg chamber despite the fact that this splice variant contains the signal peptide sequence that directs it to be secreted (Figure 3F).

### Sema2a interacts with Capu during oogenesis

The Capu co-IP and TurboID data suggested that Sema2a could be directly binding to Capu or part of a complex. To compare the subcellular localizations of Sema2a and Capu, we performed immunofluorescence experiments on ovaries expressing Capu-mScarlet-OLLAS, under control of the endogenous gene (Bailey et al., 2024). These ovaries were stained with antibodies against Sema2a and OLLAS. Consistent with previous studies, Capu localized to the follicle cells, the oocyte, and nurse cells, with enrichment at the cortex of nurse cells (Figures 4A’,B’,C’) (Bailey et al., 2024; Bor et al., 2015). In stage 8 and younger egg chambers, the Sema2a and Capu signals colocalize in the oocyte and more specifically at the posterior (Figures 4A-A’’). Like Sema2a, Capu levels in the oocyte decreased during stage 9 and their localization overlaps in the follicle cells (Figures 4B-B’’). Capu and Sema2a were also present in the migrating border cells, a cluster of 6-10 follicle cells that are necessary for the formation of the eggshell (Figure S2C, boxed in yellow) (Montell et al., 1992). By stage 11, Capu remained at the nurse cell cortex but like Sema2a, it was undetectable in the oocyte (Figure 4C-C’’). Thus, Sema2a and Capu localizations overlap in the oocyte as well as in the follicle cells, and their expression levels follow similar temporal patterns.

**Figure 4.**
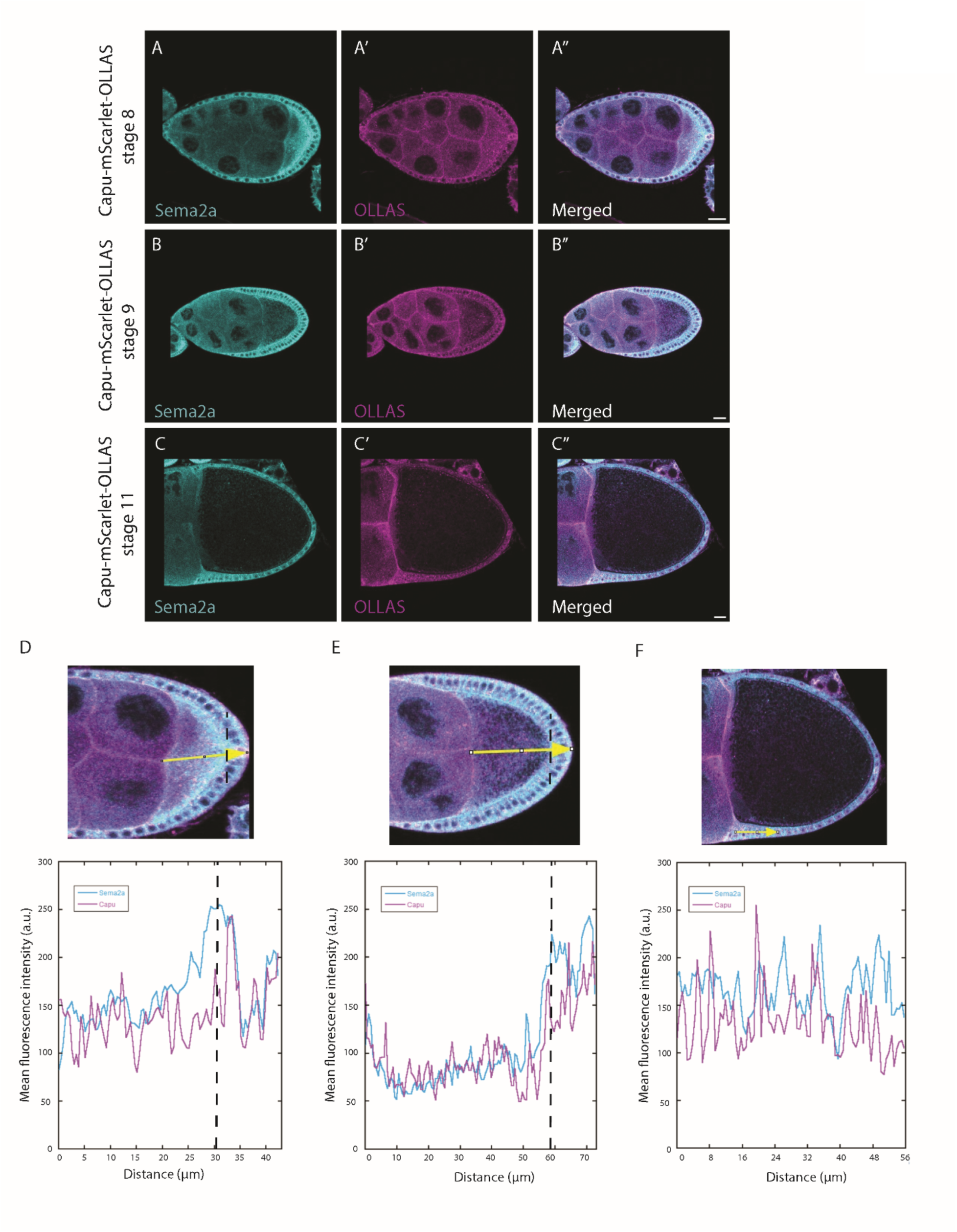
Colocalization of Sema2a and Capu. Egg chamber expressing Capu-mScarlet-OLLAS and stained with antibodies against Sema2a and OLLAS. (A) stage 8, (B) stage 9, and (C) stage 11. At stage 8, Sema2a and Capu colocalize mostly in the posterior of the oocyte and follicle cells. At stage 9, the majority of the overlap is in the follicle cells. At stage 11, Sema2a and Capu are not detected in the oocyte but remain in the follicle cells. Scale bars are 20 µm. (D-F) Intensity profiles from (A-C’’) showing similar expression patterns of Sema2a and Capu.

We further examined the localization patterns of Sema2a and Capu by analyzing intensity profiles from the anterior to the posterior of the oocyte and through the follicle cells (Figures 4D-F, yellow arrows). In stage 8 egg chambers, Sema2a and Capu filled the oocyte but the highest levels of the two were near the posterior of the oocyte and in the follicle cells (Figure 4D). In stage 9, Sema2a and Capu were most concentrated in follicle cells with the highest intensity and overlap starting at approximately the oocyte/follicle cell interface (Figure 4E). By stage 11, Sema2a and Capu were still present but not concentrated in a specific area in the follicle cells (Figure 4F).

Given that *spir-* and *capu-*null egg chambers completely lack a mesh, we examined the mesh in a Sema2a-null background. In wild-type oocytes, the mesh is present during mid-oogenesis from stages 6-10A and disappears during stage 10B. We measured mesh density by staining egg chambers with phalloidin and collected data from stage 7 and stage 9 egg chambers (Figure 5) (Dahlgaard et al., 2007; Quinlan, 2013). In each sample, we included egg chambers expressing histone-GFP, which served as a wild-type internal control to compensate for staining variability (Bor et al., 2015). In *sema2a^03021^/sema2aDf* oocytes, the mesh was not different from wild-type at stage 7 (Figures 5A and M; all statistics are given in figure legends). In contrast, mesh density decreased significantly by stage 9 (Figures 5B and M). We noted that the standard deviation at stage 7 was almost twice that at stage 9, and there seemed to be a bimodal distribution. When analyzed as two groups, one set displayed a decrease in mesh density comparable to the stage 9 level and the other, an increase in mesh density that differed significantly from each other. In fact, the mesh was difficult to detect in multiple independent runs, suggesting that it is often less dense. We note that the eggs examined were potentially biased by the issues of viability loss in Sema2a-null flies. Furthermore, a fraction of *sema2a^03021^/sema2aDf* mutants have normal-sized ovaries and may have other differences compared to the smaller ovaries. Data were collected from both ovary types but later stages usually had to be acquired from the larger ovaries.

**Figure 5.**
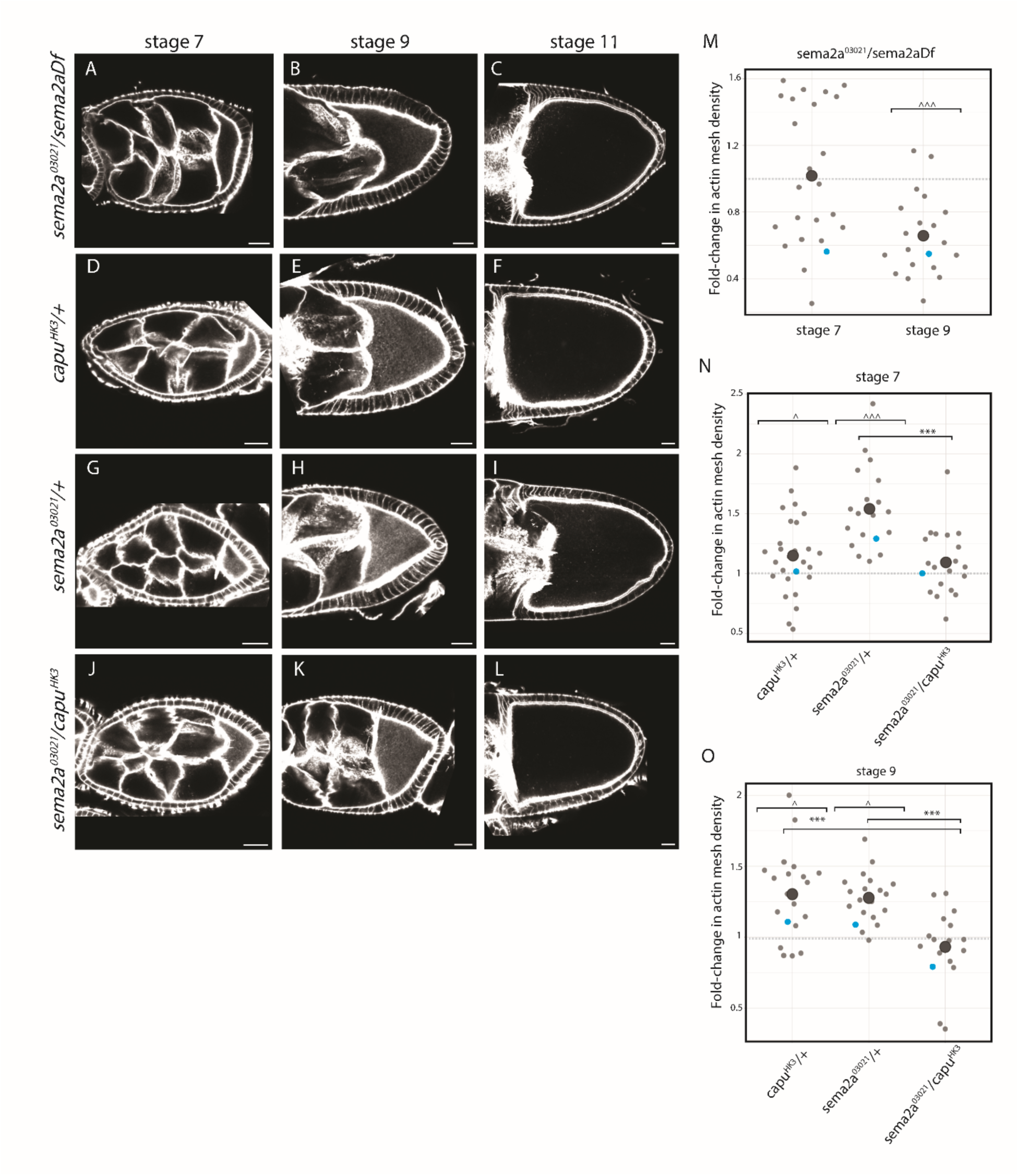
Sema2a and Capu protein levels affect actin mesh density. Egg chambers stained with phalloidin to measure actin mesh density, (A-C) *sema2a^03021^/sema2aDf,* (D-F) *capu^HK3^/+*, (G-I) *sema2a^03021^/+*, and (J-L) *sema2a^03021^*/ *capu^HK3^.* Scale bars are 20 µm. (M-O) Quantification of actin mesh density. The average intensity within the oocyte was normalized to a population of egg chambers that express Histone-GFP, stained together, as a wild-type internal control. Dotted line at 1 represents the control intensity level. Blue data points correspond to the images in (A-L). (M) Average mesh intensity is the same as wild-type in Sema2a-nulls, stage 7 (1.0 ± 0.4, n = 24, p = 0.19). At stage 9, mesh intensity is below wild-type (0.65 ± 0.24 fold-change, n = 20, p = 0.0014) (Values are means ± standard deviations and p-values were determined with a one-way ANOVA and post-hoc Dunnet’s multiple comparisons test to compare each experimental genotype against the wild-type control). When stage 7 data are analyzed as two groups, decrease in mesh density of one population is comparable to the stage 9 level (0.65 ± 0.19, n = 12), while the other had an increase in mesh density (1.38 ± 0.22, n = 12). The two groups differ significantly from each other (p < 0.001 based on a Student’s two tailed T-test performed after an F-test confirmed that the two samples have equal variance, p = 0.586). (N) Comparison of heterozygotes and transheterozygotes to wild-type reveals a genetic interaction. Stage 7 *capu^HK3^/+* mesh density is slightly higher than wild-type (1.15 ± 0.33 fold change, n = 27, p = 0.016 with Dunnet and p = 0.051 with Tukey multiple comparison tests). The mesh density of *sema2a^03021^/+* oocytes is significantly higher (1.54 ± 0.35, n = 19, p < 0.001). The stage 7 *sema2a^03021^/capu^HK3^* trans-heterozygote did not differ from wild-type (0.93 ± 0.28, n = 21, p = 0.07), consistent with a dosage compensation effect. Between samples: *capu^HK3^/+* vs. *sema2a^03021^/capu^HK3^*, p = 0.980; *sema2a^03021^/+* vs. *sema2a^03021^/capu^HK3^*, p < 0.001. (O) At stage 9, the mesh density of both heterozygotes was denser than wild-type (1.3 ± 0.2-fold change, n = 20, p = 0.031 and 1.3 ± 0.3-fold change, n = 20, p = 0.016, respectively). A one-way ANOVA and Tukey post-hoc test indicates that the two phenotypes do not differ from one another (p = 1.00). Again, the transheterozygotes did not differ from wild-type (0.9 ± 0.3, n = 17, p = 0.75). Between samples: *capu^HK3^/+* vs. *sema2a^03021^/capu^HK3^*, p < 0.001; *sema2a^03021^/+* vs. *sema2a^03021^/capu^HK3^*, p < 0.001. ***p<0.001 for comparisons between experimental samples. ^p<0.05 and ^^^p<0.001 for comparisons between experimental and Histone-GFP within same sample.

We confirmed a genetic interaction between Sema2a and Capu by crossing Sema2a (*sema2a^03021^)* and Capu (*capu^HK3^)* loss-of-function alleles to generate *sema2a^03021^/capu^HK3^* trans-heterozygotes. Viability and ovary size were not detectably affected in either the Capu heterozygote (*capu^HK3^/+*) or the trans-heterozygote (*sema2a^03021^/capu^HK3^*). However, examination of the actin mesh provided evidence of the genetic interaction. At stage 9, in both *sema2a^0302^/+* and *capu^HK3^/+* oocytes, the meshes were significantly denser than wild-type (Figures 5E,H,O). Interestingly, the mesh density in the stage 9 *sema2a^03021^/capu^HK3^*trans-heterozygote recovered; it was not different from wild-type, consistent with a dosage compensation effect (Figures 5K,O). Furthermore, both *sema2a^03021^/+* and *capu^HK3^/+* significantly differ from *sema2a^03021^/capu^HK3^*(Figure O). The same trend was apparent at stage 7 except that the increase in mesh density for *capu^HK3^/+* was less dramatic and less statistically significant (Fig 5D,G,J,N). As is the case for wild-type, no mesh was detected in stage 11 oocytes of any of the experimental genotypes (Figures 5C,F,I,L). The presence of the mesh during mid-oogenesis and its absence during late-oogenesis implies that there is a proper transition from slow to fast streaming in these genotypes. Together, the data suggest that Sema2a and Capu protein levels are important for mutual regulation and mesh assembly.

### Sema2a inhibits Capu in vitro

To determine whether Sema2a regulates Capu’s actin assembly activity, we tested purified recombinant proteins in a pyrene-actin polymerization assay. The C-terminal half of Capu, (Capu-CT, amino acids 467-1059) contains the proline-rich formin homology 1 (FH1), the well-conserved FH2, and the tail domains (Higgs and Peterson, 2005; Pruyne, 2016; Vizcarra et al., 2011). We found that Sema2a inhibited Capu-mediated actin assembly in a dose-dependent manner (4 μM actin, Figure 6A; 2 μM actin, Figure S2D). Sema2a alone had no effect on actin assembly, demonstrating that the inhibition is due to interaction between Sema2a and Capu-CT (Figure 6A).

**Figure 6.**
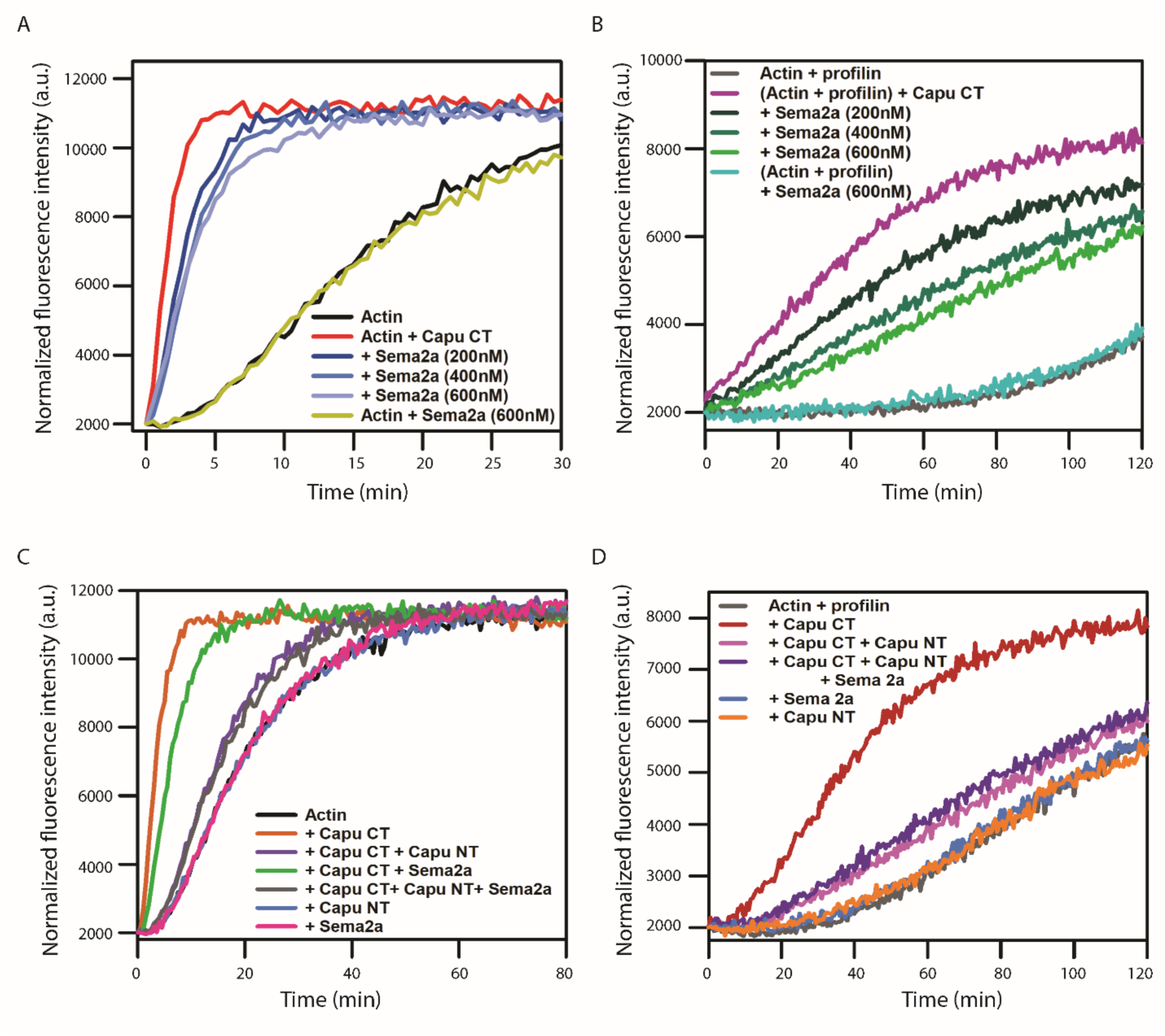
Effect of Sema2a on Capu-mediated actin assembly. (A) Representative normalized fluorescence traces of actin polymerization for samples containing 4 μM actin monomers (5% pyrene labeled), 20nM Capu CT, and a range of Sema2a. (B) Representative normalized fluorescence traces of actin polymerization for samples containing 4 μM actin monomers (5% pyrene labeled) with 8 μM human profilin, 20nM Capu CT, and a range of Sema2a. (C) Representative normalized fluorescence traces of actin polymerization for samples containing 4 μM actin monomers (5% pyrene labeled), 20nM Capu CT, 200 nM Capu NT, and 200 nM Sema2a. (D) Representative normalized fluorescence traces of actin polymerization for samples containing 4 μM actin monomers (5% pyrene labeled) with 8 μM human profilin., 20nM Capu CT, 200 nM Capu NT, and 600 nM Sema2a.

Next, we performed a similar assay in the presence of profilin, which is known to strongly inhibit Capu-CT’s nucleation activity while it accelerates elongation (Bor et al., 2012; Yoo et al., 2015). Sema2a again inhibited Capu-CT–mediated actin polymerization in a dose-dependent manner (Figure 6B). It had no impact on profilin-actin assembly.

Capu is autoinhibited by an interaction between the N-terminal Capu Inhibitory domain (CID) and the Capu-tail, C-terminal to the FH2 domain (Bor et al., 2012). We asked whether Sema2a influenced this interaction. The N-terminal half of Capu (Capu-NT, amino acids 1-466), which contains the CID, has no effect on actin assembly on its own, as we have previously reported (Figure 6C) (Bor et al., 2012). Addition of 200 nM Capu-NT reduces actin assembly in the presence of 20 nM Capu-CT to just above the baseline actin alone rate (Figure 6C). We observed no difference when Sema2a was added to this combination (Figure 6C). The same was true when profilin was present (Figure 6D). However, addition of Sema2a in the presence of a lower concentration (50 nM instead of 200 nM) of Capu-NT reduces Capu-CT-mediated actin assembly more than Capu-NT (Figure S2E). We interpret this as evidence that Capu-NT competes with Sema2a binding to Capu-CT.

Overall, these data demonstrate that Sema2a regulates Capu-CT-mediated actin nucleation directly, whether profilin is present or not. Consistent with these *in vitro* studies, the mesh density is increased in *sema2a^03021^/+* oocytes, compared to wild-type controls and *sema2a^03021^/capu^HK3^.* Together, these data suggest that lower levels of Sema2a permit higher levels of Capu activity in the oocyte (Figures 5N and O).

### Sema2a mutants disrupt osk mRNA localization

Additional evidence of interaction between Sema2a and Capu comes from examining *osk* mRNA localization in mid-oogenesis. Proper localization of polarity determinants is necessary to establish the major body axes of the oocyte and future embryo. Localization of some of these polarity determinants depends on the presence of the actin mesh during mid-oogenesis. One such component is *osk* mRNA, which is distributed throughout the ooplasm during stages 1-7 and is then directed to and enriched at the posterior pole of the oocyte from stage 8 onward (Kim-Ha et al., 1991). Other polarity factors that are localized during mid-oogenesis include *bcd* and *grk*, which localize to the anterior and anterodorsal poles of the oocyte, respectively (Thio et al., 2000; Trovisco et al., 2016). We examined the localization patterns of these mRNAs in mutant backgrounds of stages 7 and 8 oocytes. *sema2a^03021^/sema2aDf* mutants showed normal localization of *bcd* and *grk,* comparable to that of wild-type (*w*^1118^) at all stages (Figures S3A-H’). In contrast, there was major disruption of *osk* mRNA in *sema2a^03021^/sema2aDf* oocytes, evidenced by clumps or foci as opposed to the more diffuse distribution of *osk* mRNA in wild-type and *capu^HK3^/+* oocytes (Figures 7A-A’’, D-D’’, G-G’’, J-J’’, M-M’’). While a fraction of *sema2a^03021^/+* oocytes (40%, n=25) showed clumping of *osk* mRNA, the phenotype was highly penetrant in *sema2a^03021^/capu^HK3^* (100%, n=14) and *sema2a^03021^/sema2aDf* egg chambers (97% n=31) (Figure 7P). This coalescence of clumps was detectable as early as stage 3 (Figures S3I-I’’). We observed comparable abnormal foci of Staufen (Stau) in *sema2a^03021^/sema2aDf* mutants and *sema2a^03021^/capu^HK3^* trans-heterozygotes, consistent with Stau’s role in *osk* mRNA transport (Figures 6C,F,I,L,O) (Micklem et al., 2000). There was no evidence of premature Osk translation (Figures 7B,E,H,K) (Besse et al., 2009; Kim-Ha et al., 1995). By stage 9, there was some evidence of residual mRNP aggregation in *sema2a^03021^/capu^HK3^* oocytes, but the majority of *osk* mRNA and Stau were at the posterior pole, and Oskar translation appeared wild-type (Figures S4A-O).

**Figure 7.**
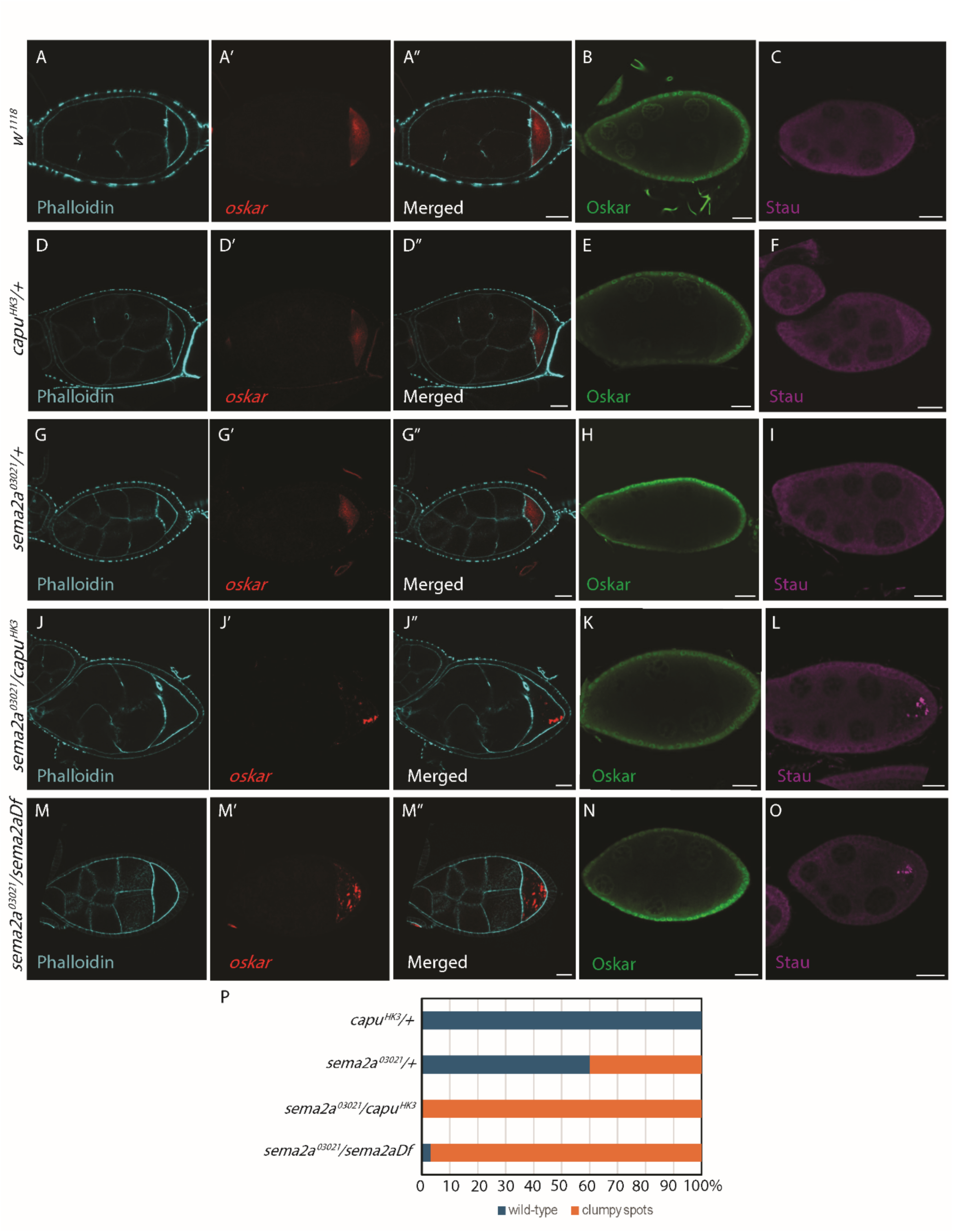
Sema2a/Capu trans-heterozygote and Sema2a-nulls disrupt *osk* mRNA localization. Stage 7 egg chambers stained with phalloidin and *osk* mRNA probe, (A-A’’) *w^1118^*, (D-D’’) *capu^HK3^/+*, (G-G’’) *sema2a^03021^/+*, (J-J’’) *sema2a^03021^*/ *capu^HK3^*, and (M-M’’) *sema2a^03021^/sema2aDf.* Both *sema2a^03021^*/ *capu^HK3^*and *sema2a^03021^/sema2aDf* show displacement of *osk* to concentrated puncta, or clumps, compared to wild-type and heterozygotes. Scale bars are 20 µm. (B, E, H, K, N) Oskar protein staining for the respective constructs. Translation and localization of Osk is not impacted in any genotype. Scale bars are 20 µm. (C, F, I, L, O) Stau protein staining for the respective constructs. Stau displacement matches that of *osk* in *sema2a^03021^*/ *capu^HK3^* and *sema2a^03021^/sema2aDf* egg chambers. Scale bars are 20 µm. (P) Quantification of different phenotypes for *capu^HK3^/+* (n=18), *sema2a^03021^/+* (n=25), *sema2a^03021^*/*capu^HK3^*(n=14), and *sema2a^03021^/sema2aDf* (n=31). *sema2a^03021^*/ *capu^HK3^* and *sema2a^03021^/sema2aDf* have all or almost all clumpy spots compared to the heterozygotes.

Similar clumps of *osk* mRNA are not observed in *spir-* or *capu*-nulls. Instead, *osk* mRNA appears scattered and fails to localize to the posterior pole at stage 9, presumably due to absence of the mesh and presence of premature fast streaming (Dahlgaard et al., 2007; Manseau and Schupbach, 1989; Theurkauf, 1994; Wellington et al., 1999). Given that *osk* mRNA clumps are present in *sema2a^03021^/capu^HK3^* at stage 7 but mesh density is indistinguishable from wild-type in the trans-heterozygote, the mislocalization of *osk* is unlikely to be a direct result of altered mesh. Instead, the difference in phenotypes suggests that Sema2a and Capu play more than one role, directly or indirectly, in mRNA localization.

### Sema2a and its canonical receptor PlexB have different phenotypes

Because the major Sema2a isoform identified was the canonical isoform A that has been extensively studied in the CNS, we chose to also examine its interacting partner, PlexB (Ayoob et al., 2006). Recall that PlexB was present in the Capu-TurboID dataset but not at significant levels (FC ≈ 3.5, p = 0.173). We tried to generate PlexB-nulls using the loss-of-function allele *plexB^KG00878^*. However, we did not recover any homozygous *plexB^KG00878^* flies (Ayoob et al. 2006 recovered ∼6% escapers). Instead, when we crossed *plexB^KG00878^* to a deficiency line (Df(4)M101-62f) (*plexB^KG00878^/plexB-Df)*, we recovered 5 or fewer escapers from each cross (Ayoob et al., 2006 recovered none). The *plexB^KG00878^/plexB-Df* mutants all survived CO_2_ exposure, with viability and ovary size resembling that of wild-type, unlike *sema2a^03021^/sema2aDf* mutants. In assessing the *osk* mRNA localization in stage 7, we found that *plexB^KG00878^/+* heterozygotes had multiple phenotypes – 50% with diffuse mRNA localization similar to wild-type, 39% with clumpy spots similar to *sema2a^03021^/sema2aDf,* and 11% (n=36) with a distinct single spot in the center of the oocyte (Figures 8A-A’’, P). The coalescence of one single spot of *osk* is comparable to the phenotypes seen in stage 9 Par-1 and *didum* mutants among others (Doerflinger et al., 2006; Krauss et al., 2009). These proportions shifted in *plexB^KG00878^/plexB-Df* egg chambers, such that only 5% had apparently normal mRNA localization, 28% had clumpy spots, and the majority, 67%, contained a single cluster of *osk* mRNA in the center of the oocyte (n=18) (Figure 8D-D’’, P). As expected, the Stau distribution in *plexB^KG00878^/plexB-Df* oocytes tracked that of *osk* mRNA in the same background (Figure 8F). However, *plexB^KG00878^/+* heterozygotes predominantly contained Stau foci, inconsistent with the majority of *osk* mRNA localization being wild-type and more comparable to that of *sema2a^03021^/sema2aDf* (Figure 8C). By stage 9 *osk* mRNA and Stau were concentrated at the posterior pole, and Oskar translation appeared wild-type, in both *plexB^KG00878^/+* and *plexB^KG00878^/plexB-Df* oocytes (Figures S5A-F).

**Figure 8.**
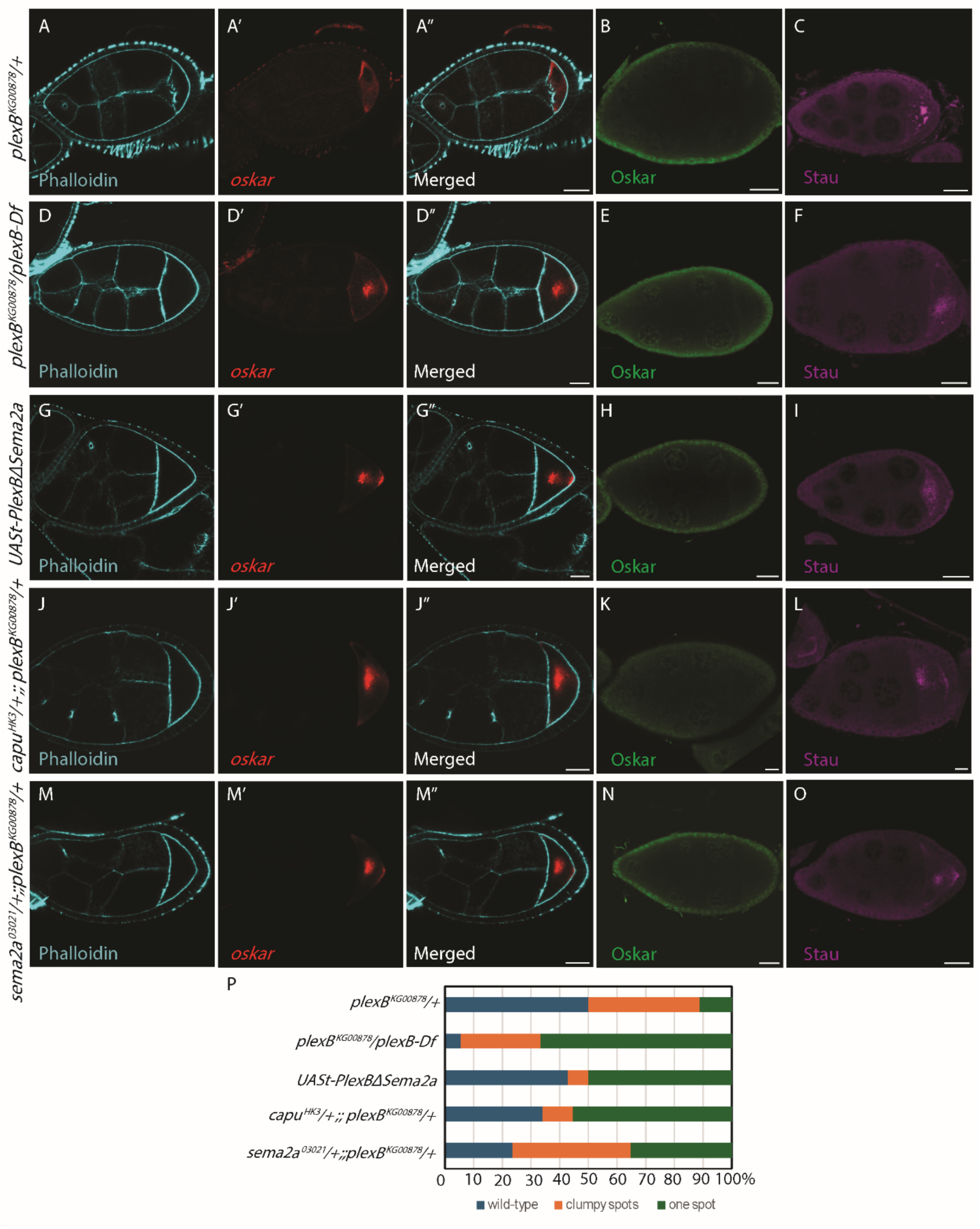
PlexB mutants exhibit *osk* mRNA localization that is different from Sema2a mutants. Stage 7 egg chambers stained with phalloidin and *osk* mRNA probe. Phenotypes varied between diffuse, one large spot, and clumps of *osk* mRNA. (A-A’’) *osk* mRNA is diffuse in most *plexB^KG00878^/+* chambers. A large single spot of osk mRNA is observed in most (D-D’’) *plexB^KG00878^/plexB-Df*, (G-G’’) UASt-PlexBΔSema2a, and (J-J’’) *capu^HK3^*;; *plexB^KG00878^* oocytes. Similar frequency of clumpy and single spot *osk* mRNA in (M-M’’) *sema2a^03021^/ plexB^KG00878^*. Scale bars are 20 µm. (B, E, H, K, N) Oskar protein staining for the respective genotypes. Translation and localization of Osk is not impacted in all genotypes. Scale bars are 20 µm. (C, F, I, L, O) Stau protein staining for the respective genotypes. Central cluster or spots of Stau matches that of *osk*. Scale bars are 20 µm. (P) Quantification of different phenotypes for *plexB^KG00878^/+* (n=36), *plexB^KG00878^/plexB-Df* (n=18), UASt-PlexBΔSema2a (n=14), *capu^HK3^*;; *plexB^KG00878^*(n=47), *sema2a^03021^/plexB^KG00878^* (n=17). Most PlexB mutants have one central spot unlike Sema2a mutants which only had clumpy spots.

To test for a role that PlexB might play in the Sema2a/Capu pathway, we crossed *plexB^KG00878^* to *sema2a^03021^* or *capu^HK3^* to generate trans-heterozygotes. We also drove expression of PlexB lacking the Sema2a binding domain (UASt-PlexBΔSema2a) using the matα-GAL4 driver. We frequently observed one central concentration of *osk* mRNA in stage 7 *capu^HK3^/+;;plexB^KG00878^/+* (55%, n=47), UASt-PlexBΔSema2a (50%, n=14), and *sema2a^03021^/+;;plexB^KG00878^/+* oocytes (35%, n=17) (Figures 8G-G’’, J-J’’, M-M’’, P). We note that *sema2a^03021^/+;; plexB^KG00878^/+* oocytes had 41% clumpy spots, consistent with some overlap in the pathways they influence. Of the 11% that contained more concentrated spots in *capu^HK3^/+;;plexB^KG00878^/+*oocytes, the *osk* mRNA distributions seemed to capture transient images of mRNA being moved toward to the posterior pole, akin to multiple transport defects (Figures S5P-P’’). In all of these backgrounds, the Stau distribution tracked the *osk* mRNA (Figures 8I, L, O) and there was no evidence of premature or ectopic Osk protein translation (Figures 8B,E,H,K,N). Once again, by stage 9, *osk* mRNA, Osk protein, and Stau were all properly localized to the posterior pole of the oocyte in these different *plexB* mutant backgrounds (Figures S5A-O). The altered *osk* mRNA localization patterns suggest that Sema2a and PlexB both operate in the *osk* transport pathway. The predominance of centralized *osk* mRNA rather than clumps and the wild-type ovary size in *plexB* mutant backgrounds suggest that Sema2a plays a role that is independent of PlexB in oogenesis.

## Discussion

### Defining the Spir/Capu interactome using complementary proteomic approaches

Combining both co-immunoprecipitation and TurboID-proximity labeling enables us to capture both transient and stable interactions, potentially offering a wide range of spatial and temporal information. We discovered proteins that were consistent with previous studies, such as Myosin V in both Spir co-IPs and TurboID datasets, suggesting that it is a stable and spatially close interaction (Figure 1B’, 2B, Table 1, Table 2). On the other hand, Spir was not enriched in Capu co-IPs but was enriched in Capu-TurboID datasets, indicating that it is spatially close but perhaps too weak or transient of an interaction to be detected in co-IPs (Figure 1C, 2C, Table 1, Table 2). This notion fits with protein localization studies and a model of Spir/Capu interaction, whereby Spir and Capu coordinate to nucleate actin, then releasing Capu to elongate the filaments independently (Bailey et al., 2024; Bradley et al., 2019; Quinlan, 2013). We note again that Capu was not expected in either Spir co-IP or Spir TurboID datasets because the SpirC isoform does not contain the KIND domain that binds to Capu.

It was intriguing to find that multiple neuronal-related proteins were enriched in Capu co-IPs, such as Sema2a, Sema2b, and homer. In fact, homologs of Capu, Fmn2 in mammals and Fmn2b in zebrafish, play roles in neurons (Almuqbil et al., 2013; Kundu et al., 2020; Nagar et al., 2021). Other parallels between the central nervous system and reproductive system have been reported. For example, Vap33 (VAMP-associated protein) is an essential gene critical to synaptic homeostasis (Kamiyama et al., 2025). It was also identified in Egl-TurboID datasets from ovaries as well as shown to be enriched in oocytes and transported in a Egl/Dynein-dependent manner (Baker et al., 2021). In addition, several proteins critical for transport and post-transcriptional regulation of mRNAs in the ovary, such as Nanos and Pumilio, play important roles in dendrite morphogenesis (Ye et al., 2004).

We performed secondary screens on a number of proteins enriched in the co-IP and TurboID datasets. While not yet independently validated, we expect that proteins found highly enriched in both co-IPs and TurboID are likely to be bona fide interactors. Preliminary data suggest that several will provide new opportunities for understanding Spir-Capu regulation and mesh dynamics.

### Sema2a plays a role in Drosophila oogenesis

We report here a previously unidentified role for Sema2a in the female reproductive system. What is most unusual is the intracellular localization of the protein. TurboID detection, immunofluorescence, and stage-specific changes in enrichment argue strongly that Sema2a is acting within the oocyte. Another semaphorin, Sema5c is known to operate in the follicle cells of the developing egg chamber (Stedden et al., 2019). In contrast to Sema2a, Sema5c acts through its canonical signaling pathway with PlexA. The protein is planar-polarized and reduces “backwards” motility, coordinating essential epithelial migration. Sema5c functions as a transmembrane protein. It is also detected intracellularly in a sparse and punctate pattern, consistent with endosomes and regulation of transmembrane protein signaling.

Interestingly, human semaphorins, including the ortholog of Sema2a, SEMA3A, play multiple roles in female fertility (Emery et al., 2023; Giacobini, 2015; Zhang et al., 2025). In fact, SEMA3A levels increase in the plasma of women with depleted ovary reserves. Thus, SEMA3A may be an effective biomarker for diagnosis and treatment (Palese et al., 2024). To date, the known ovarian roles of semaphorins are classical signaling roles, with the help of plexin and neuropilin receptors. They influence follicle development and ovulation, both remotely through the gonadotropin hormone system and by locally influencing cell motility and remodeling (Messina and Giacobini, 2013; Zhang et al., 2025).

*Drosophila* Sema2a is clearly critical to oogenesis. Absence of Sema2a has a strong impact on ovary development, resulting in a majority of ovaries that are considerably smaller than wild-type, even not detected at times (Figure 3B-C). Development appears to stall during early to mid-oogenesis in these small ovaries. The presence of normal-sized ovaries that proceed to later stages in the Sema2a-nulls suggests that the ovary may compensate for the loss of Sema2a to some degree, which could contribute to the complexity we observe in mesh density data.

Inhibition of Capu-mediated actin assembly is consistent with some of the mesh phenotypes we observed. Higher mesh intensity correlated with decreased levels of functional Sema2a in *sema2a^03021^/+* oocytes. In addition, in the *sema2a^03021^/capu^HK3^* trans-heterozygote mutants, the mesh density is similar to that of wild-type, suggesting that relative amounts of these two proteins are important for function. However, the mesh in the Sema2a-null was usually less dense than wild-type. Perhaps, alternate regulatory pathways overcompensate in the absence of Sema2a.

The CID and Spir-KIND both inhibit Capu by binding to the Capu-tail. Based on the lack of an effect when Sema2a was added to Capu-CT plus Capu-NT, we speculate that Sema2a binds in the same region. However, Sema2a is not a particularly potent Capu-inhibitor. The K_i_ is hundreds of nanomolar, whereas Ki’s for Capu-NT and the Spir-KIND domain are ∼10 nM (Bor et al., 2012; Vizcarra et al., 2011). The fact that we identified Sema2a in Capu co-IPs, suggests that the *in vivo* interaction is fairly stable. Together, these data suggest that the stability of the interaction may depend on a complex of proteins. Notably, Sema2a functions as a dimer and can heterodimerize with Sema2b, which was also enriched in the Capu interactome.

Previous investigations showed no correlation between mesh density and streaming velocities beyond the all or nothing impact of the *capu*-null (Bor et al., 2015; Emmons et al., 1995; Yoo et al., 2015). Furthermore, egg chambers that lack a mesh continue to develop well past mid-oogenesis. Together, these data lead us to conclude that mesh density is not the critical factor impacting development when it fails in Sema2a mutants during mid-oogenesis. The fact that a strong *osk* mRNA phenotype in the same *sema2a^03021^/capu^HK3^* transheterozygote that rescues the mesh phenotype indicates that at least two pathways are mediated by Sema2a-Capu interaction.

### *osk* mRNA localization

The mechanisms controlling *osk* mRNA are dependent on the interplay of many different and complex processes, including the cooperation of the cytoskeleton, molecular motors, and repression/activation factors that contribute to proper localization of *osk* mRNA and its spatiotemporally controlled translation (Lasko, 2012; Lehmann, 2016). During stages 7 and 8, microtubules are oriented with plus-ends biased toward the posterior of the oocyte, enabling the plus-end directed motor, kinesin, to transport *osk* mRNA across the oocyte (Brendza et al., 2000; Parton et al., 2011; Zimyanin et al., 2008). During this time, *osk* mRNA is silenced by Bruno1 and other proteins to prevent ectopic translation of Osk protein (Besse et al., 2009; Kim-Ha et al., 1995; Mhlanga et al., 2009). By stages 9-10, Bruno1 repression is lifted and Makorin1 is able to stabilize and anchor Osk (Dold et al., 2020).

The consequences of *osk* mislocalization specifically during stage 7 and earlier are unknown and need to be further investigated. The clumps of concentrated *osk* mRNA that are formed in *sema2a^03021^/capu^HK3^*and *sema2a^03021^/sema2aDf* mutants are similar to those detected in PTB (polypyrimidine tract-binding protein) mutants (Figure 7) (Besse et al., 2009). PTB is part of the ribonucleoprotein complex required for *osk* mRNA transport, which contributes to translation suppression. Consistently, premature Osk protein colocalizes with the abnormal *osk* mRNA punctae in the PTB-mutant oocyte. The fact that Osk protein does not appear ectopically in Sema2a mutants in stages 8 and earlier and properly translates during stage 9 and later suggests that the Bruno1 and Makorin1 pathway is functioning.

The accumulation of *osk* mRNA foci rather than the typical diffuse distribution throughout the oocyte could be due to a cytoskeletal malfunction. Dosage compensation in *sema2a^03021^/capu^HK3^*transheterozygotes that results in wild-type actin mesh density during stage 7 oocytes is in direct contrast with the severe *osk* mRNA phenotype observed at this stage. This observation suggests that *osk* mRNA mislocalization is not a consequence of an altered actin mesh. The fact that *bcd* and *grk* are properly localized at the anterior suggests that microtubules and their motors are not entirely dysfunctional. However, bcd and grk are localized at the pointed ends of microtubules, which may be normally distributed even when the plus ends are not.

We found one other report of aggregated *osk* mRNA that was corrected by stage 9. Snee and MacDonald (Snee and Macdonald, 2009) found that sponge bodies can be organized along a spectrum from a diffuse distribution to a more clumpy, reticulated structure. They found that virgin females and flies that are not fed extra yeast, in addition to the typically rich food they grow on, were more likely to have reticulated sponge bodies. In contrast, 3-5 day old females fed a normal diet supplemented with dry yeast had consistently diffuse small sponge bodies. In the initial report, *osk* mRNA and Stau localized to the clumps but not consistently. Based on FRAP data and the fact that *osk* mRNA and Stau were properly concentrated at the oocyte posterior by stage 9, it was concluded that the interaction between RNPs and the sponge body are transient as opposed to structural or functional. Our experiments were performed under conditions that favor diffuse sponge bodies. Perhaps Sema2a and Capu play a role in sponge body organization, resulting in clumps in these mutants.

## Conclusions

The discovery of interaction between Capu and Sema2a is an excellent example of the power of unbiased proteomics approaches. Neither would be on a list of predicted interactors with the other, based on their known roles and localizations, i.e. intracellular vs. secreted. Nevertheless, the genetic, cell biological, and biochemical data strongly support the proteomics data. Notably, Sema2b was also enriched in our proteomics datasets. Sema2a functions as a dimer and can heterodimerize with Sema2b (Rozbesky et al., 2019). In fact, there is evidence of significant crosstalk between the Sema2 proteins and the Plexin receptors (Winberg et al., 1998; Wu et al., 2011). It remains to be seen if Sema2b functions with Sema2a in oogenesis. It will be important to determine if Sema2b upregulation is responsible for compensation in the Sema2a-null background. It is curious that Sema2a’s canonical receptor PlexB also function in a pathway that temporarily alters *osk* localization when disrupted yet gives a distinct phenotype at the whole animal level, the ovary level, and in the specific mRNA mislocalization. Many exciting questions remain, including how Sema2a is retained intracellularly, whether it plays additional intracellular roles, and how the transient mislocalization of *osk* affects other processes in oocyte progression, embryogenesis, and overall fertility.

## Methods

### Drosophila stocks

The following fly lines were obtained from Bloomington Drosophila Stock Center: *w*^1118^, Sema2a^03021^/SM5 (RRID:BDSC_11257, (Kolodkin et al., 1993)), Df(2R)B65/CyO (RRID:BDSC_65746, (Wu et al., 2011)), PlexB^KG00878^/ciD (RRID:BDSC_14579 (Bellen et al., 2004)), PlexB-Df(4)M101-62f/eyD (RRID:BDSC_9433), UASp-SpirC-GFP (RRID:BDSC_24765 (Rosales-Nieves et al., 2006)), His2AV-EGFP/SM6A (RRID:BDSC_24163). UASp-CapuA-GFP, UASp-GFP-CapuA, Capu-mScarlet-OLLAS, and capu-GAL4 were constructed in the Quinlan lab and reported in earlier publications (Bailey et al., 2024; Quinlan, 2013). *capu*^HK3^ was provided by the Schupbach lab (Manseau and Schupbach, 1989) and UASp-GBP-TurboID, matα-GAL4 was built and generously provided by the Gonsalvez lab (Baker et al., 2021).

### Co-immunoprecipitation

Flies expressing UASp-CapuA-GFP/capu-GAL4 or UASp-SpirC-GFP/capu-GAL4 were fattened with yeast paste overnight at 25°C. 40-100 pairs of ovaries were dissected in 1X PBS and homogenized in lysis buffer (50 mM Tris-HCl pH 7.5, 150 mM NaCl, 1 mM EDTA, 1% Nonidet P40 Substitute, protease inhibitor cocktail). The homogenized samples were sonicated at 40% amplitude with 1 second pulses for 30 seconds. Lysates were spun down and the resulting supernatants were incubated with 25 μL of equilibrated GFP-Trap magnetic agarose (ChromoTek) bead slurry for an hour at 4°C. For experiments with Latrunculin A, 10 µM was added. After an hour, unbound proteins were removed with wash buffer (50 mM Tris-HCl pH 7.5, 150 mM NaCl, 0.5 mM EDTA, 0.05% Nonidet P40 Substitute, protease inhibitor cocktail). Bound proteins were eluted with 200 mM glycine pH 2.5 and immediately neutralized with 1 M Tris-Cl pH 10.4. Eluted proteins were either run on a gel and analyzed by Western blot or prepped for mass spectrometry. Proteomics experiments were analyzed in triplicate from three biologically independent runs.

### Mass spectrometry

The eluted proteins were reduced with 5 mM Tris (2-Carboxyethyl) phosphine (TCEP) and alkylated with iodoacetamide (IAA) at RT for 30 min in the dark. SP3 protein clean-up was performed using carboxylate-modified magnetic beads at a ratio of 10:1 (w/w) beads to protein (Hughes et al., 2019). Protein binding was facilitated using ethanol (EtOH) to a final concentration of 50% and washed 3 times with 80% EtOH. Bound beads were then resuspended in 2 M urea and 100 mM Tris-HCl pH 7.5 and digested with 1:100 (w/w) LysC and 1:25 (w/w) trypsin overnight at 37°C, mixing at 1200 rpm. The next day, peptide binding was facilitated using 100% acetonitrile (ACN) and washed 3 times with 100% ACN. Peptides were eluted with 2% dimethyl sulfoxide (DMSO) at 37°C for 30 min. The eluate was collected, dried, and resuspended in 5% trifluoroacetic acid (TFA) for LC-MS/MS analysis.

Samples were analyzed with data-independent acquisition (DIA)-based quantitative proteomics using an UHPLC system coupled to an Orbitrap Astral mass spectrometer. Peptides were loaded onto a Pepsep C18 reverse-phase column (15 cm x 150 µm) at 52°C and electrosprayed into an Orbitrap Astral mass spectrometer. Peptide separation was performed using the Vanquish Neo UHPLC system, with mobile phase A and B consisting of 0.1% formic acid (FA) in water and 0.1% FA in ACN, respectively. A 15-minute gradient was used as follows: mobile phase B was increased to 5% from 0-1 min, 5-15% from 1-5 min, 15-25% from 5-12.6 min, 25-38% from 12.6-13.6 min, 38-80% from 13.6-13.7 min, and remained at 80% from 13.7-15 min. DIA spectra were acquired in positive ion mode with MS1 parameters as follows: resolution of 240,000, m/z range of 380-980, normalized AGC target of 500%, and maximum injection time of 3 ms. MS2 spectra were collected at 80,000 resolution, HCD energy of 25%, normalized AGC target of 500%, and maximum injection time of 7 ms. Thermo RAW files were processed using DIA-NN with peptide lengths set to 7-30 and maximum number of missed cleavages as 2. An *in silico* spectral library was generated from the *Drosophila melanogaster* reference proteome.

### Proximity labeling

Flies expressing UASp-GBP-TurboID, matα-GAL4/UASp-CapuA-GFP or UASp-GBP-TurboID, matα-GAL4/UASp-SpirC-GFP were fattened with yeast paste overnight at 25°C. 50 pairs of ovaries were dissected in 1X PBS and homogenized in lysis buffer (50 mM Tris-HCl pH 7.5, 150 mM NaCl, 1 mM EDTA, 1% Nonidet P40 Substitute, protease inhibitor cocktail). Homogenized samples were sonicated at 40% amplitude with 1 second pulses for 30 seconds. Lysates were spun down and the resulting supernatant was incubated with 10 μL of equilibrated streptavidin magnetic bead (ThermoFisher) slurry for two hours at 4°C. Unbound proteins were washed 3x with lysis buffer, 3x with 1% SDS, 3x lysis buffer, 3x high salt lysis buffer, 3x lysis buffer, and 4x with PBS (Baker et al., 2021). After extensive washes, proteins were reduced, alkylated, and digested on beads overnight at 37°C, mixing at 1200 rpm. The next day, SP3 clean-up was performed starting from the ACN step. Proteomics experiments were analyzed in triplicate.

### Antibodies

The following antibodies were used: chicken anti-GFP (Aves Labs Inc.; 1:1000), mouse anti-Sema2a (Developmental Studies Hybridoma, 1:20), rat anti-OLLAS (Novus Biologicals; 1:2500), rabbit anti-Osk (gift from Ephrussi lab; 1:3000), mouse anti-Stau (gift from Doe lab, 1:50). The following secondary antibodies from Jackson Immunologicals were all used at 1:200: donkey anti-chicken AlexaFluor488, donkey anti-mouse AlexaFluor647, donkey anti-rat AlexaFluor488, donkey anti-rabbit AlexaFluor488. Samples were counterstained with rhodamine phalloidin (1:600), AlexaFluor647 phalloidin (1:600), or streptavidin AlexFluor647 (1:5000) (Molecular Probes).

### Immunofluorescence

Flies were fattened with yeast paste overnight at 25°C. Samples were fixed using previously published protocols (Bailey et al., 2024; Cooley et al., 1992; Ong et al., n.d.). Briefly, ovaries were dissected in cold 1X PBS and fixed in 5% PFA, 16% 1X PBS, and hexane. After 10 minutes of fixation, ovaries were washed 3 times with PBST (1X PBS, 0.1% Triton X-100, 0.5% BSA). Ovaries were then stained with primary antibodies overnight at 4°C. The next day, ovaries were washed with PBST 3 times. The appropriate secondary antibodies were added for 2 hours at room temperature and then ovaries were washed again 3 times with PBST. Ovaries were mounted in ProLong Gold and imaged the next day. Quantification of colocalization was performed using Fiji. Each channel of the image was normalized and a segment of interest was drawn (width: 5 and size: 10). Plot profile was used to generate a graph of intensity vs. distance (of the drawn segment).

### Actin mesh staining

Samples were fixed using previously published protocols (Dahlgaard et al., 2007; Quinlan, 2013). Briefly, ovaries were dissected in 10% PFA in 1X PBS. Ovaries were washed 4 times with PBST and stained with 1 µM phalloidin for no longer than half an hour. Ovaries were washed again 4 times and mounted in ProLong Gold for imaging the next day. Quantification of the density of the actin mesh was performed using Fiji. The mean intensity of each oocyte was compared to the mean intensity of histone-GFP expressing ovaries stained in the same tube to calculate the fold-change of each fly construct (e.g. *sema2a^03021^/capu^HK3^*mean intensity/histone-GFP mean intensity). Data were presented using SuperPlotsOfData (Goedhart, 2021). Statistical analysis of mesh density was performed by one-way ANOVA and post-hoc Dunnet and Tukey multiple comparison tests. Means ± standard deviations and adjusted p-values are reported in the figure legends.

### smiFISH

Samples were prepared according to previously published protocols (Calvo et al., 2021; Lu et al., 2023; Tsanov et al., 2016). The probes were described in (Bailey et al., 2024). Briefly, ovaries were dissected in 1X PBS and washed with PBST and smiFISH wash buffer (2X solution of 0.3 M sodium citrate and 3 M sodium chloride (SSC), 10% deionized formamide). Then, the appropriate annealed mRNA probes were added in hybridization buffer (10% dextran sulphate, 2X SSC, 10% deionized formamide) overnight at 37°C. The next day, ovaries were washed with smiFISH wash buffer and PBST and mounted in ProLong Gold for imaging the following day.

### Microscopy

Localization, mesh, and smFISH images were acquired using Zeiss LSM 700 confocal microscope with a 40x/1.3 or 63x/1.4 oil objective (Banerjee lab). Brightfield images of whole ovaries were acquired using Zeiss Axio Zoom.V16 Macro Zoom with X-Cite TURBO LED 6-Channel light source, driven by SlideBook (3i).

### RT-PCR

*w*^1118^ flies were fattened with yeast paste overnight at 25°C. 100 pairs of ovaries were dissected in 1X PBS. Total RNA extraction was performed as described (Bremer et al., 2024). Sema2a cDNA was synthesized with SuperScript III using a gene-specific primer 5’ CTTGAATTTGCCATTGAAGGCAGC 3’. Ethanol precipitation was performed and the purity and concentration of the cDNA was assessed with a NanoDrop. The cDNA was amplified with Phusion High-Fidelity DNA Polymerase using the following steps: initial denaturation 98°C, 0:30; 30 cycles 98°C, 0:30, 68.2°C, 0:30, 72°C, 0:30, final extension 72°C, 5:00, and infinite hold 4°C.

For sequence present in all isoforms:

Forward primer: 5’ GAACGAAGATCGAGATACGCTCTATG 3’

Reverse primer: 5’ CTTGAATTTGCCATTGAAGGCAGC 3’

For Sema2a isoform A:

Forward primer: 5’ CCATTTGCCAGTGAAATCAATTCC 3’

Reverse primer: 5’ TAAATCGCAGTTGAGTTGTCGAGG 3’

The amplified product was sent to Plasmidsaurus to confirm sequence.

Detailed thermocycler protocols and primers for Sema2a isoforms C and E are available upon request.

### Protein expression and purification

*Acanthamoeba castellani* actin, Capu-CT, and Capu-NT constructs were expressed and purified using published protocols (Vizcarra 2011, Bor et al 2012). Sema2a protein was gifted by Daniel Rozbesky.

Briefly, Capu-CT and Capu-NT were expressed in Rosetta competent cells and grown in Terrific Broth at 37°C until they reached an OD_600_ of 0.6-0.7. Cells were induced with 0.25 mM isopropyl-β-d-thiogalactoside at 18°C overnight. The resulting cell pellets were harvested, flash frozen with liquid nitrogen, and stored at -80°C. Purification of Capu-CT was performed using Talon resin and MonoQ anion exchange as described (Vizcarra et al., 2011). Purification of Capu-NT was performed using glutathione agarose resin and the protein was cleaved from GST using PreScission Protease overnight at 4°C. For CapuNT, the protein was concentrated and gel-filtered on the Superdex 200 10/300 GL column as described (Bor et al., 2012). Purified fractions were dialyzed into storage buffer (20 mM Tris pH 8.0, 100 mM NaCl, 1 mM DTT, 50% glycerol). Aliquots were flash frozen with liquid nitrogen and stored at -80°C. The concentration of the Capu-CT construct was calculated based on quantitative Sypro-Red (Invitrogen) staining while the concentration of Capu-NT was based on the extinction coefficient from Bor et al. (Bor et al., 2012) Human Profilin-1 was expressed and purified as described for Drosophila profilin (Chic) (Bor et al., 2012). Profilin-1 concentration was determined using the extinction coefficient 14,992 M−1 cm−1.

### Pyrene-actin polymerization assays

Pyrene assays were performed essentially as described (Bor et al., 2012). Actin polymerization was initiated by the addition of 2× concentrated protein mix in 2× polymerization buffer (1× KMEH buffer: 10 mM HEPES, pH 7.5, 1 mM EGTA, 1 mM MgCl_2_, 50 mM KCl, and 0.2 mM ATP) to 50 μl of 2× concentrated Mg^2+^- ATP-actin at a final concentration of 4 μM with 5% pyrene-labeled actin. Mg^2+^- ATP-actin was prepared just before initiation of the assay by incubating Ca^2+^-actin in G-buffer with ME exchange buffer (final concentration, 0.05 mM MgCl_2_ and 0.2 mM EGTA) for 2 minutes at room temperature. The final concentrations of additional proteins used are indicated in figures legends. For reactions containing profilin, actin was preincubated human profilin (at twice the actin molar ratio) prior to use. Pyrene fluorescence was monitored for 1.5-2 hours (with excitation at 365 nm and emission at 407 nm) at 25°C, in an Infinite M Nano+ plate reader (Tecan). The time between mixing of components and the beginning of data collection ranged between 20-30 seconds for each assay.

## Acknowledgements

We thank Graydon Gonsalvez for providing us with the TurboID fly line and Daniel Rozbesky for the purified recombinant Sema2a protein. We also thank members of the Quinlan and Wohlschlegel lab for their help and support. Thanks to Orkun Akin for providing many reagents and useful discussions as well as Emil Reisler for lending resources. C.W. especially thanks family and friends for their love and support. This work was supported by NIH grant R01GM096133 to M.E.Q., R35GM153408 to J.W., and award number T32GM145388 to C.W.

---

**Figure S1.**
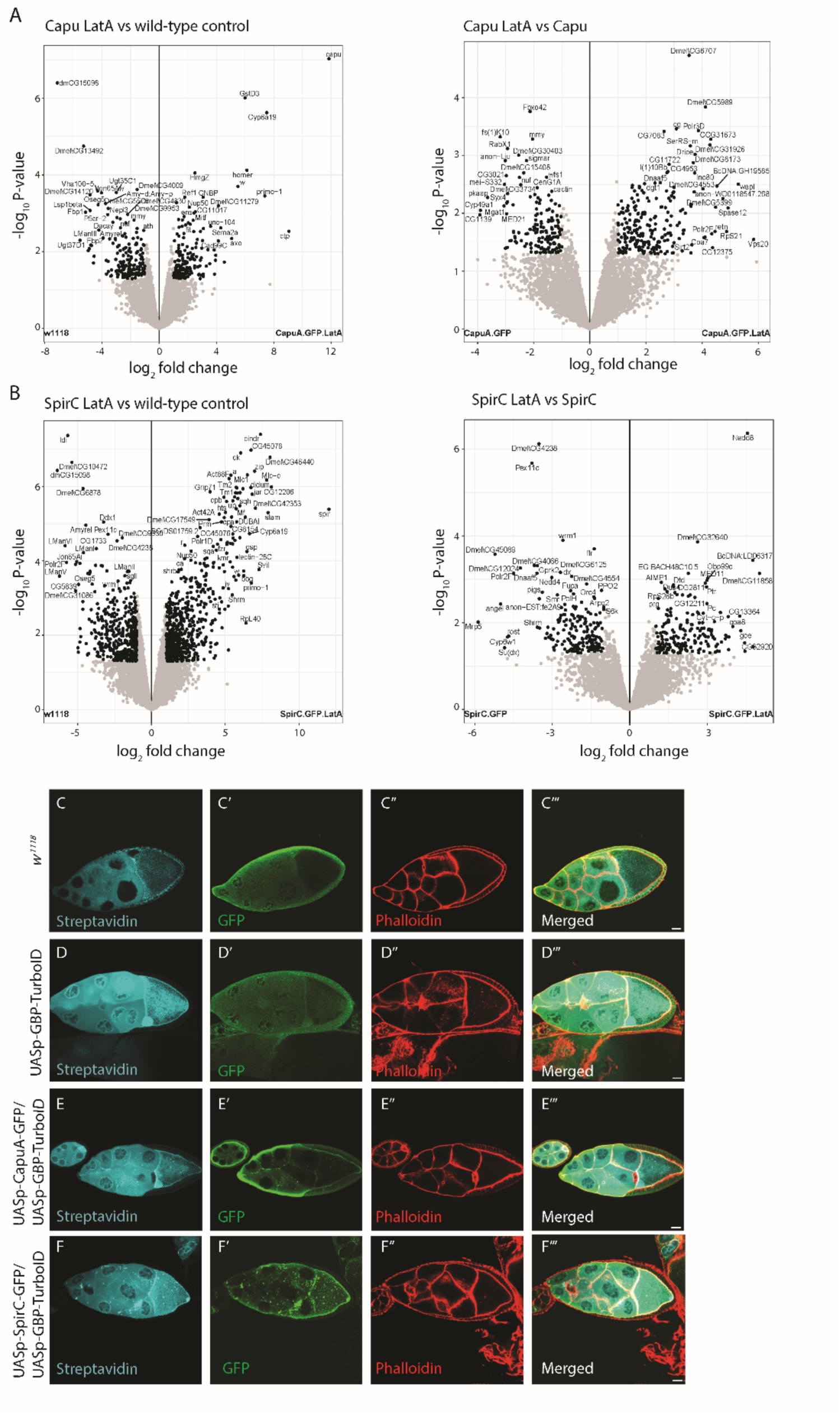
Enrichment of co-IPs performed with Latrunculin A present and validation of proximity labeling. (A) Volcano plots showing Capu with addition of 10 µM Lat A compared to wild-type or compared to Capu without addition of LatA. First plot shows same proteins – Ctp, Sema2a, Homer, etc. Second plot shows that there is no change between ± Lat A. (B) Volcano plots showing Spir with addition of 10 µM Lat A compared to wild-type or compared to Spir without addition of LatA. First plot shows same proteins – didum, Cpb, Cindr, etc. Second plot shows that there is no change between ± Lat A. Egg chambers stained with streptavidin, anti-GFP, or phalloidin (C-C’’’) *w^1118^,* (D-D’’’) UASp-GBP-TurboID driven by matα-gal4, (E-E’’’) UASp-GBPTurboID and UASp-CapuA-GFP driven by matα-gal4, and (F-F’’’) UASp-TurboID and UASp-SpirC-GFP driven by matα-gal4. The colocalization of GFP-tagged Capu and Spir and streptavidin signal suggests successful biotinylation of proximal proteins within ∼10 nm radius of Capu and Spir. Scale bars are 20 µm.

**Figure S2.**
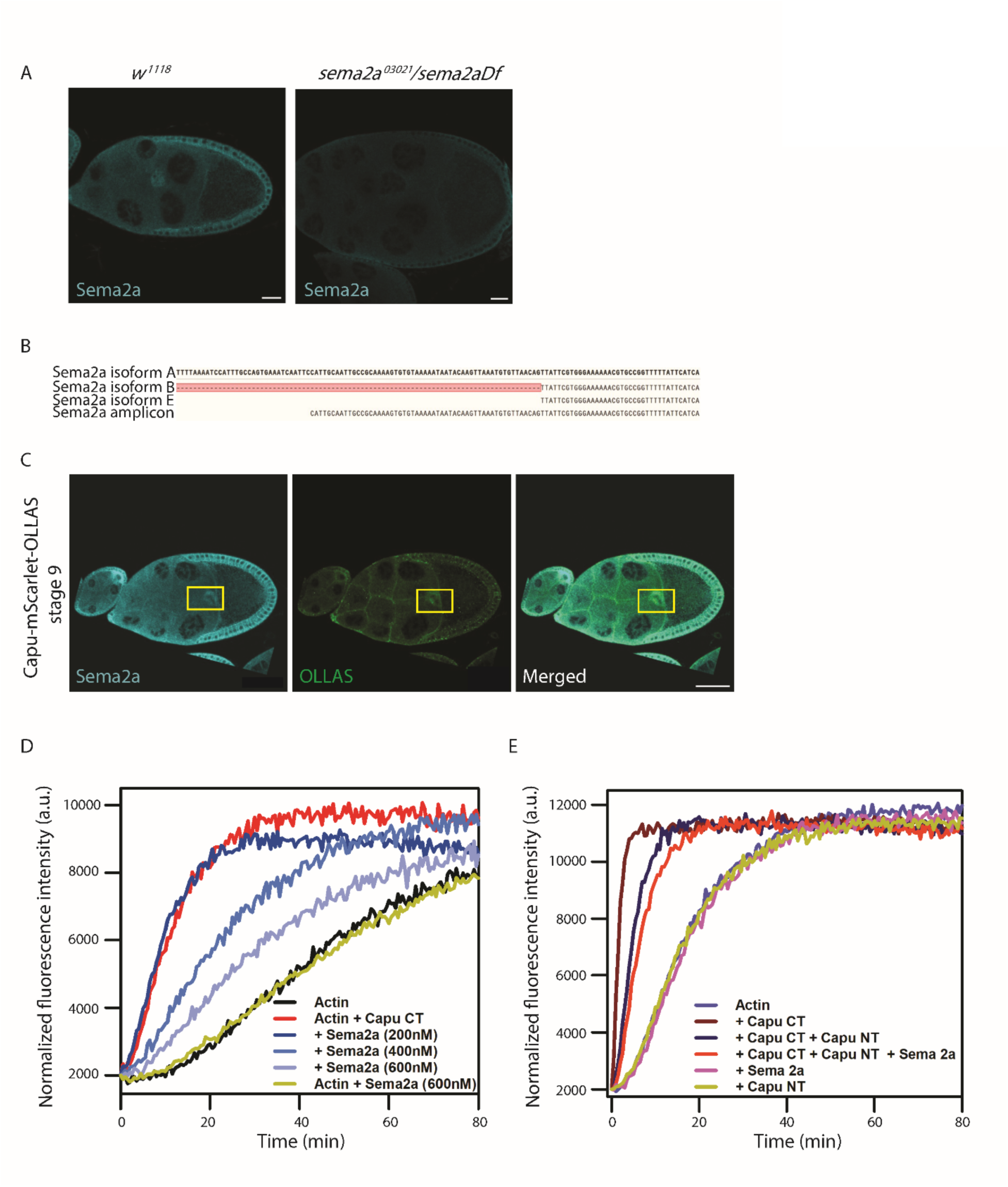
Sema2a localization, isoform, and interaction with Capu. (A) Sema2a staining for *w^1118^* and *sema2a^03021^/sema2aDf* stage 9 egg chambers. *w^1118^* shows abundant signal in egg chamber compared to *sema2a^03021^/sema2aDf.* Scale bars are 20 µm. (B) Plasmidsaurus sequencing data comparing Sema2a amplicon to Sema2a isoform sequences. Sema2a amplicon matches that of Sema2a isoform A. (C) Capu-mScarlet-OLLAS egg chambers stained with Sema2a and OLLAS show signal in the migrating border cells, boxed in yellow. Aside from the well-known role that Capu plays in the mesh, this observation could indicate other unexplored roles of both Capu and Sema2a in border cells. Scale bars are 50 µm. (D) Representative normalized fluorescence traces of actin polymerization for samples containing 2 μM actin monomers (5% pyrene labeled), 20nM Capu CT, and a range of Sema2a. (E) Representative normalized fluorescence traces of actin polymerization for samples containing 4 μM actin monomers (5% pyrene labeled), 20nM Capu CT, 50 nM Capu NT, and 200 nM Sema2a.

**Figure S3.**
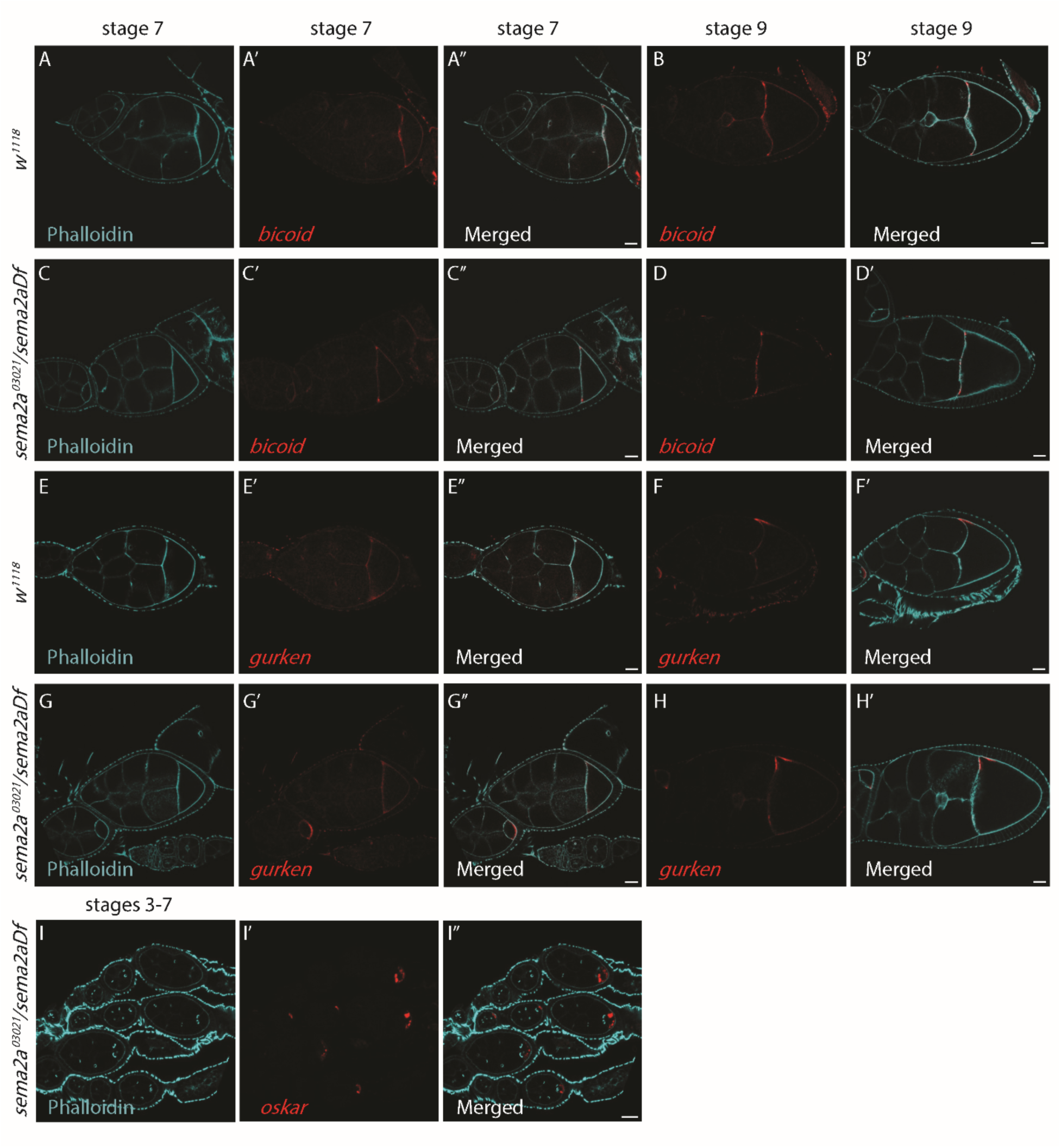
*bicoid, gurken, and oskar* mRNA localization in *sema2a^03021^/sema2aDf* oocytes. Stage 7/8 or stage 9 egg chambers probing for *bicoid (bcd)* (A-B’) *w^1118^* and (C-D’) *sema2a^03021^/sema2aDf.* Localization of *bcd* is normal and comparable to that of wild-type. Stage 7/8 or stage 9 egg chambers probing for *gurken (grk)* (E-F’) *w^1118^*and (G-H’) *sema2a^03021^/sema2aDf.* Localization of *grk* is also unaffected and is similar to that of wild-type. Scale bars are 20 µm. (I-I’’) Stages 3-7 egg chambers of *sema2a^03021^/sema2aDf* stained with phalloidin and *osk* mRNA probe. Disruption of *osk* mRNA starts in very early stages of oogenesis, beginning as early as stage 3.

**Figure S4.**
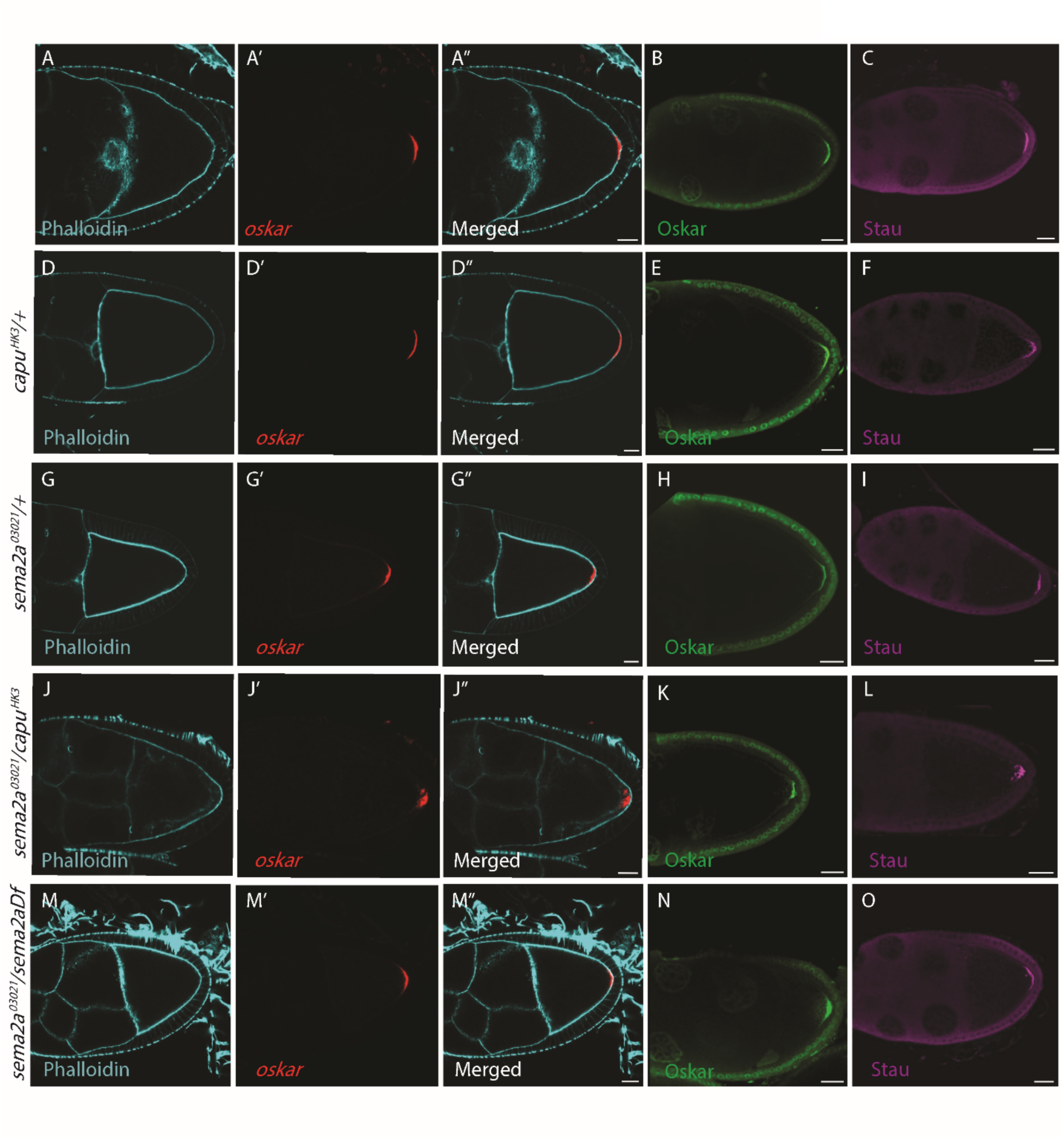
Sema2a/Capu trans-heterozygotes and Sema2a-nulls have normal *osk* mRNA localization by stage 9. Stage 9 egg chambers stained with phalloidin and *osk* mRNA probe, (A-A’’) *w^1118^*, (D-D’’) *capu^HK3/+^*, (G-G’’) *sema2a^03021^/+*, (J-J’’) *sema2a^03021^*/ *capu^HK3^*, and (M-M’’) *sema2a^03021^/sema2aDf.* The *sema2a^03021^*/ *capu^HK3^* egg chambers have some residual RNP aggregation and clumps while *sema2a^03021^/sema2aDf* have normal *osk* distribution compared to wild-type and heterozygotes. Scale bars are 20 µm. (B, E, H, K, N) Oskar protein staining for the respective constructs. Translation and localization of Osk is normal and matches that of *osk* mRNA. Scale bars are 20 µm. (C, F, I, L, O) Stau protein staining for the respective constructs. Stau localization is also normal in all genotypes. Scale bars are 20 µm.

**Figure S5.**
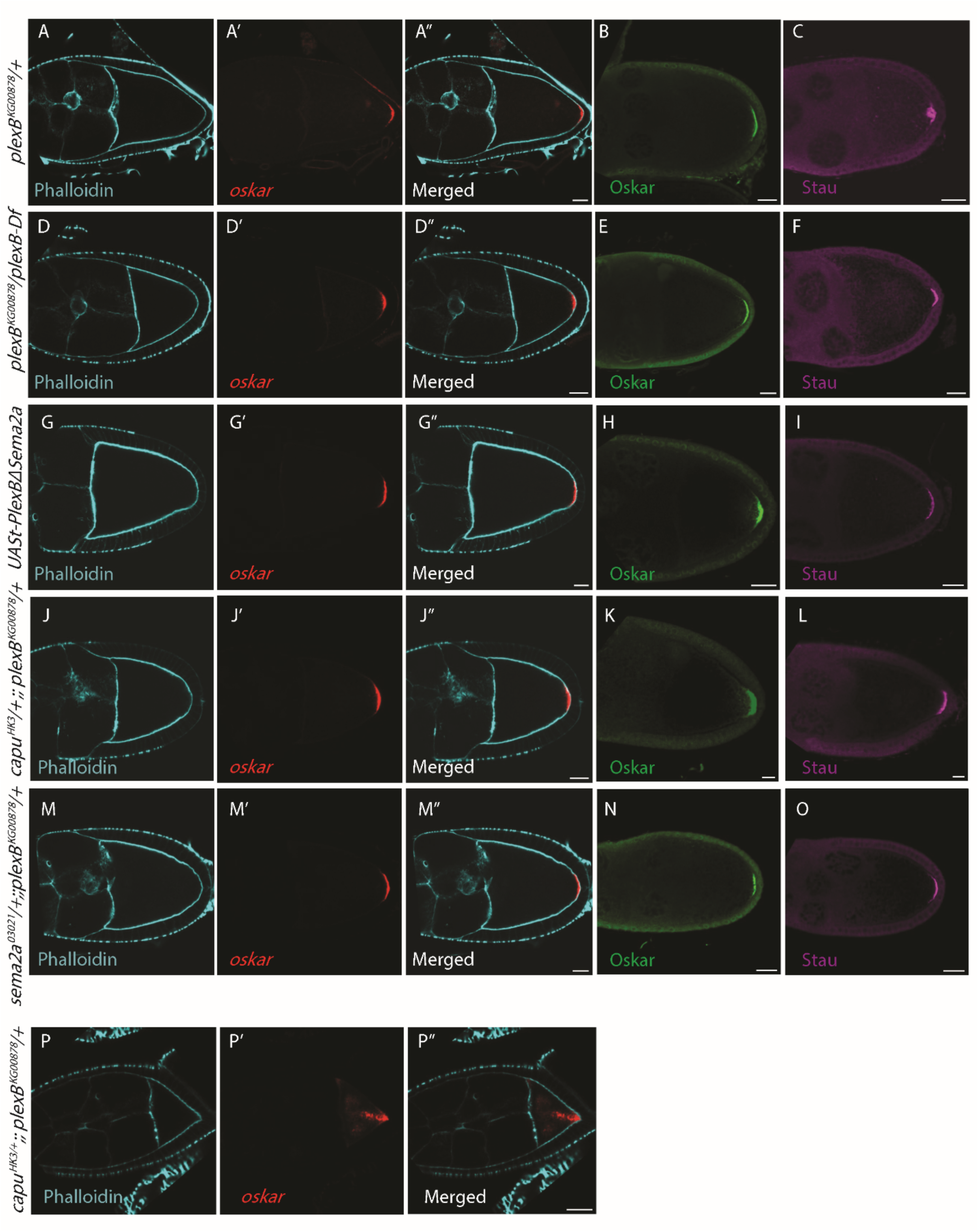
PlexB mutants display *osk* mRNA localization that is normal by stage 9. Stage 9 egg chambers stained with phalloidin and *osk* mRNA probe. (A-A’’) *plexB^KG00878^/+* chambers, (D-D’’) *plexB^KG00878^/plexB-Df*, (G-G’’) UASt-PlexBΔSema2a, (J-J’’) *capu^HK3^*;; *plexB^KG00878^* oocytes, and (M-M’’) *sema2a^03021^/ plexB^KG00878^*. *osk* mRNA localization is localized at the posterior and comparable to wild-type in all genotypes. Scale bars are 20 µm. (B, E, H, K, N) Oskar protein staining for the respective genotypes. Translation and localization of Osk is again normal in all genotypes shown. Scale bars are 20 µm. (C, F, I, L, O) Stau protein staining for the respective genotypes. Stau is also normal, similar to both *osk* mRNA and Osk protein. Scale bars are 20 µm. (P-P’’) Another phenotype seen in the PlexB-Capu trans-heterozygotes. Stage 8 egg chamber stained with phalloidin and probed for *osk* mRNA. Localization of *osk* shows clumpy spots being transported to the posterior of the oocyte, suggesting defects in transport mechanism. Scale bars are 20 µm.

